# Epigenetic restoration of differentiation competency via reversal of epiblast regionalisation

**DOI:** 10.1101/2024.12.27.630149

**Authors:** Magdalena A. Sutcliffe, Steven W. Wingett, Charles A.J. Morris, Eugenia Wong, Stefan Schoenfelder, Madeline A. Lancaster

## Abstract

Although the epiblast in the embryo has the capacity to generate all tissues of the body, its *in vitro* counterparts often exhibit differentiation biases, posing significant challenges for both basic research and translational applications involving pluripotent stem cells (PSCs). The origins of these biases remain incompletely understood. In this study, we identify PSC differentiation biases as arising from fluctuations in repressive and activating histone posttranslational modifications, leading to the acquisition of a caudal epiblast-like phenotype.

We present a novel approach to overcome this bias using a chemical chromatin restoration (CHR) treatment. This method restores transcriptional programs, chromatin accessibility, histone modification profiles, and differentiation potential, effectively recapitulating the competent anterior epiblast-like state. Furthermore, we propose that a high bivalency state is a defining feature of the anterior human epiblast. We suggest that fluctuations in histone modification marks drive epiblast regionalization, ultimately shaping cellular responses to differentiation cues.

## Introduction

In the process of embryonic development a single cell generates the remarkably broad repertoire of tissues that build a fully formed body, and amongst them the most complex organ – the brain. In the primate embryo, after the inner cell mass of the blastocyst separates into epi- and hypoblast, the epiblast generates the amniotic cavity, and acquires a transient pluripotent state poised for generation of all three germ layers.^1^ The posterior part of the embryo is exposed to WNT signals that initiate primitive streak formation, whereas the periphery of the embryonic disc perceives BMP ligands from the amnion with which it is continuous.^2–4^ The anterior part of the epiblast, however, is shielded from those patterning signals, and acquires the default fate of neural ectoderm.^5–7^

When epiblast equivalent cells are cultured *in vitro* without any patterning factors they can form cerebral organoids, or, when presented with appropriate developmental cues, they can generate all other cell types of the body.^8,9^ However, both human embryonic stem cells (ESCs) and induced pluripotent stem cells (iPSCs) derived from individual donors differ markedly in their capacity to differentiate into desired cell types *in vitro*.^10–17^ The molecular underpinnings of this functional human PSC heterogeneity remain poorly understood, and this has been a major hurdle in PSC-based disease modelling approaches and regenerative cell therapies.

Hypotheses to explain inter-individual PSC differentiation heterogeneity have mainly focused on genetic variation (both naturally occurring and acquired), and changes in epigenetic marks, such as DNA methylation, as the primary drivers of differences in human PSC potential.^14,18,19^ However, accumulating evidence also shows that the precise developmental stage and state at which the differentiation cues are received plays a crucial role in defining human PSC differentiation outcomes.^1,20^

Studies of mouse embryos demonstrated that even before pluripotency exit, epiblast cells show signs of regionalisation, and although all cells from mouse epiblast are still pluripotent, they show differentiation bias.^21^ Interestingly, these features are preserved *in vitro*: mouse epiblast cells cultured in a dish show features of caudalisation and are incompatible with anterior neural fate unless Wnt signalling is inhibited.^22–24^ However, if mouse cells are cultured from naïve conditions equivalent to the pre-implantation stage and then allowed to proceed to primed pluripotency *in vitro*, they can acquire either anterior neural fate or posterior neuromesodermal identities upon pluripotency exit.^25^ This suggest that primed pluripotency is competent for antero-posterior patterning and that the A-P axis is established before germ layer specification.^26^ However, whether regionalisation is mutually exclusive with pluripotency, how it affects the interpretation of developmental cues, and how the earliest mechanisms establish regional identity during early development remain to be elucidated.

In this study, we leverage the natural diversity between human PSC lines to investigate the mechanism of fate decisions around the peri-implantation epiblast regionalisation point to ask how this process underlies cell competency to generate anterior neural (ANE) tissues. Since the timing of cell fate transitions is preserved *in vitro*, we take advantage of the protracted transition from blastocyst to gastrulation in the human embryo to capture more subtle and gradual transition states.^27–29^ Free of constraints and compensatory mechanisms present in natural embryos, and with hugely expanded analytical tools we can probe and dissect their precise identities.^30^ We demonstrate that low ANE competency is associated with a posterior epiblast-like state, which at the molecular level is characterised by a locus-specific erosion of bivalent chromatin marks that leads to the aberrant expression of key developmental transcription factor genes. We further show that high chromatin bivalency is crucial for ANE specification and efficient development of other germ layers. Notably, we find that changes in bivalent chromatin at differentially expressed gene loci are largely uncoupled from changes in chromatin accessibility or DNA methylation.

Further, we establish a defined protocol that restores full PSC differentiation capacity in non-competent PSC lines by resetting their aberrant transcription and chromatin states. We predict that this approach will become instrumental for a broad range of PSC-based applications in developmental biology, disease modelling and regenerative medicine. Finally, we highlight the power of leveraging inter-individual genetic and epigenetic variation across the human population to decipher the signalling pathways and gene regulatory circuits that govern cell lineage specification during early human development.

## Results

### Transcriptomic differences underlying competency

To study ANE competency we selected a model of unguided cerebral organoids that faithfully recapitulates forebrain development from pluripotent stem cells (PSCs). In this model, PSCs are aggregated into embryoid bodies, exit pluripotency upon withdrawal of TGFβ and FGF2 and self-organise into tissues closely resembling foetal dorsal telencephalon.^9^ The cerebral organoid method is particularly sensitive to the quality of the starting PSC line and the differentiation outcomes are only partially improved by optimising PSC culture conditions.^31^ To elucidate underlying features of PSCs that predict cerebral organoid generation success, or competency, we examined the transcriptomic landscape of a range of PSC lines. To ensure detection of cell-intrinsic differences, rather than differences due to culture conditions or other extrinsic sources of variation, we re-analysed single-cell (sc)RNA-seq data from cerebral organoids generated previously in our laboratory from a pool of human induced PSCs from healthy donors^16^ in which the cells were cultured in the same dish before forming organoids. Within these mixed culture organoids, individual cell lines produced varying proportions of neural and non-neural identities (Figure 1A).

**Figure 1.**
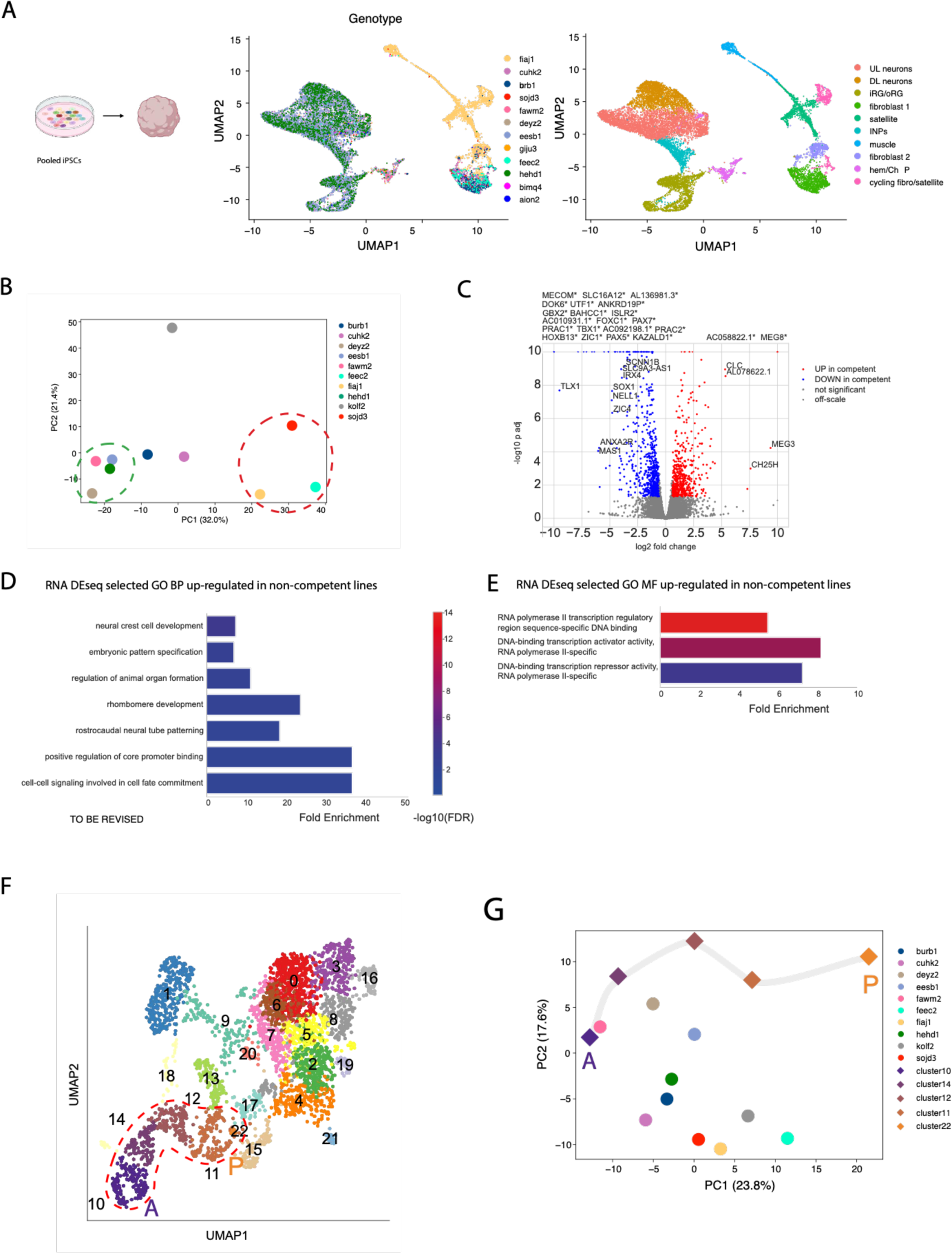
A - uniform manifold approximation and projections (UMAPs) of the pooled iPSC-derived cerebral organoids; left UMAP summarizes the cell line genotype distribution and right UMAP summarizes the organoid cell types, B - scatter plot of the first two principal components of the principal competent analysis of bulk transcriptomes of selected iPSC lines (differential genes defined by VST threshold), encircled in green are the lines unambiguously classified as competent, in red the lines unambiguously classified as non-competent, C – volcano plot of differentially expressed genes between competent (eesb1, hehd1, kolf2) and non-competent (fiaj1, sojd3, feec2) lines, D – bar plot of selected significantly enriched terms from Panther gene ontology (GO) Biological Process (BP) analysis of transcripts up-regulated in non-competent lines, E - bar plot of selected significantly enriched terms from Panther GO Molecular Function (MF) analysis of transcripts up-regulated in non-competent lines, F – UMAP summarizes cell populations in day 8 human stem cell embryo model, encircled are the epiblast clusters 10, 14, 12, 11 and 22, G – scatterplot of the first two principal components of the principal competent analysis of the bulk transcriptomes of selected iPSC lines together with “pseudo-bulked” epiblast clusters of the day 8 SEM (analysis of 250 top variable genes).

From these data, we were able to identify PSC lines that showed a clear ability to properly generate brain cells (deyz2, eesb1, fawm2, hehd1), demonstrating they were competent, while others generated mainly non-brain identities (burb1, cuhk2, feec2, fiaj1, sojd3), indicating they were non-competent (Figure 1S A, B). We focused on cell lines with the highest proportion of cells represented in the scRNA-seq data, and analysed available bulk RNA-seq datasets for the PSC lines themselves to test for differences already present in the pluripotent state. To include a male karyotype not already present among the competent lines in our sample, we added data from kolf2 cells, previously demonstrated to be competent.^31^ PCA analysis revealed separation of competent and non-competent lines that was most pronounced in the first principal component (Figure 1B), and evident in the dendrogram (Figure 1S C).

This demonstrated that transcriptomic differences between competent and non-competent cell lines already exist in the starting PSC culture. To identify the particular genes and processes within PSC lines that contribute to competency, we performed DEseq analysis on 6 lines that showed the clearest phenotypes: 3 competent lines (eesb1, hehd1, kolf2) and 3 non-competent lines (feec2, fiaj1 and sojd3) (Figure 1C, Table S1). GO term analysis of genes upregulated in non-competent lines returned several biological process (BP) terms associated with embryonic patterning and differentiation, especially of the posterior structures (Figure 1D, Table S1). Molecular function (MF) analysis pointed toward transcriptional regulators and in particular developmental transcription factors (Figure 1E).

We speculated that the aberrant expression of transcripts associated with body development might reflect a regional bias of the non-competent lines similar to the primitive streak bias of mouse EPiSCs.^32^ To test this possibility, we turned to data from human embryos and stem cell embryo models (SEM). SEM data offer the possibility to capture stages of development when material would otherwise be difficult to access and where very limited cells are represented.^33^ In particular, we were interested in examining the pre-gastrulation epiblast when the primitive streak anlage is first established. We cross-referenced the day 8 SEM data with an integrated scRNA-seq dataset of human embryos and a spatial transcriptome of human gastrula for cluster annotation.^3,34^ We identified 22 clusters using Leiden resolution of 1.2 that captured regionalisation of the epiblast cluster into two anterior subclusters (clusters 10 and 14) and three posterior subclusters (clusters 11, 12 and 22, Figure 1F, Figure 1S D). To identify parallels between PSC lines and different regions of epiblast we created pseudo-bulk data objects from these epiblast clusters 10, 11, 12, 14 and 22. We then performed dimensionality reduction analysis of the top 250 most variable genes. PC1 of our analysis captured the anterior to posterior gradient of the SEM data (Figure 1G), and at the same time separated competent and non-competent lines, with the exception of kolf2. PC2 reflected the separation of PSC and SEM data. These data support a more caudal character of the non-competent lines.

### Elevated endogenous WNT signalling in non-competent lines

*In vivo*, the establishment of the anterior-posterior (A-P) axis is instructed by a Wnt signalling gradient, which in mouse occurs about one day before the appearance of the primitive streak. ^35,36^ *In vitro* experiments further confirmed that Wnt is sufficient for A-P regionalization and that the regionalisation occurs prior to germ layer specification.^25,26,37^ Because of the instructive role of Wnt in caudalising the embryo, we hypothesised that differences in competent and non-competent PSC lines might be due to differences in WNT activity.

Although our culture conditions were devoid of WNT ligands, endogenous WNT activity has been reported in human PSCs and was demonstrated to confound differentiation outcomes, in particular restricting anterior fate acquisition.^24,38,39^ To explore endogenous WNT signalling, we analysed expression of WNT ligands in our cultures (Figure 1S E, Table S2). WNT transcripts were generally expressed at very low levels but canonical WNT3 was higher in three of the non-competent lines and in the kolf2 line. Non-canonical WNT5A and B were expressed in all lines but at the highest levels in non-competent fiaj1 and sojd3 lines. Since A-P patterning results from canonical WNT signalling, we then explored expression of canonical WNT targets previously reported in human ES cells.^35,40^ We detected upregulation of all WNT-responsive transcripts in the non-competent feec2 line and higher abundance of HHEX, MIXL1 and GSC in the sojd3 and fiaj1 lines (Figure 1S F). Surprisingly, the competent kolf2 line also showed some degree of upregulation of WNT targets but in a pattern distinct to non-competent cells.

To test whether the endogenous WNT signalling in non-competent lines was the source of the posterior bias, we treated two of the non-competent lines, burb1 and sojd3, with the pan-WNT inhibitor IWP2 for 8 passages (4 weeks) while culturing on laminin 521 in Stemflex, and then generated cerebral organoids from them. We analysed the obtained tissues by immunofluorescence on days 10 and 20. One of the cell lines, burb1, showed improvement in the identity and morphology of the organoids (Figure 1S G). The sojd3 line, however, did not show any improvement (Figure 1S H), demonstrating that while spontaneous WNT activity may underly non-competence for some cell lines, its inhibition is not sufficient to restore competency more generally.

### Chromatin accessibility differences between competent and non-competent lines

We next considered the possibility that the failure of WNT inhibition to restore competence may be due to an irreversible state of the cells, perhaps in the form of cellular memory.^41,42^ Exposure to WNT signal produces cellular memory that changes differentiation outcomes in micropatterned hPSC colonies.^43,44^ We therefore hypothesised that a similar phenomenon could be responsible for the regionalisation of the non-competent cell lines, and that in the case of the IWP2 non-responsive line sojd3, the memory of WNT pre-exposure could not be erased simply by WNT inhibition.

The establishment and propagation of cellular memory often involves epigenetic changes at the chromatin level.^45^ To investigate chromatin differences between competent and non-competent lines that could potentially explain the resistance of non-competent cells to WNT inhibition and other culture modifications,^31^ we explored global chromatin accessibility. We performed ATAC-seq analysis of 4 non-competent lines and 4 competent lines. In general, chromatin accessibility was similar between all samples and almost identical in regions within +/- 5kb of transcription start sites (Figure 2A), indicating no differences in global chromatin accessibility. However, differential peaks revealed common trends among the non-competent compared with competent lines. Principal component analysis showed clustering of all 4 competent lines and 3 non-competent lines separately when 220,897 variable peaks were considered (Figure 2B-C). Interestingly, the burb1 cell line clustered with the competent lines, perhaps reflecting its status as a line that is not irreversibly non-competent, as also demonstrated by its repair with WNT inhibition. However, because of this, it was omitted from the subsequent differentially accessible peak analysis. The patterns of differentially accessible peaks in competent and non-competent lines also confirmed that burb1 cells were more similar to competent cells than non-competent cells (Figures 2D-E).

**Figure 2.**
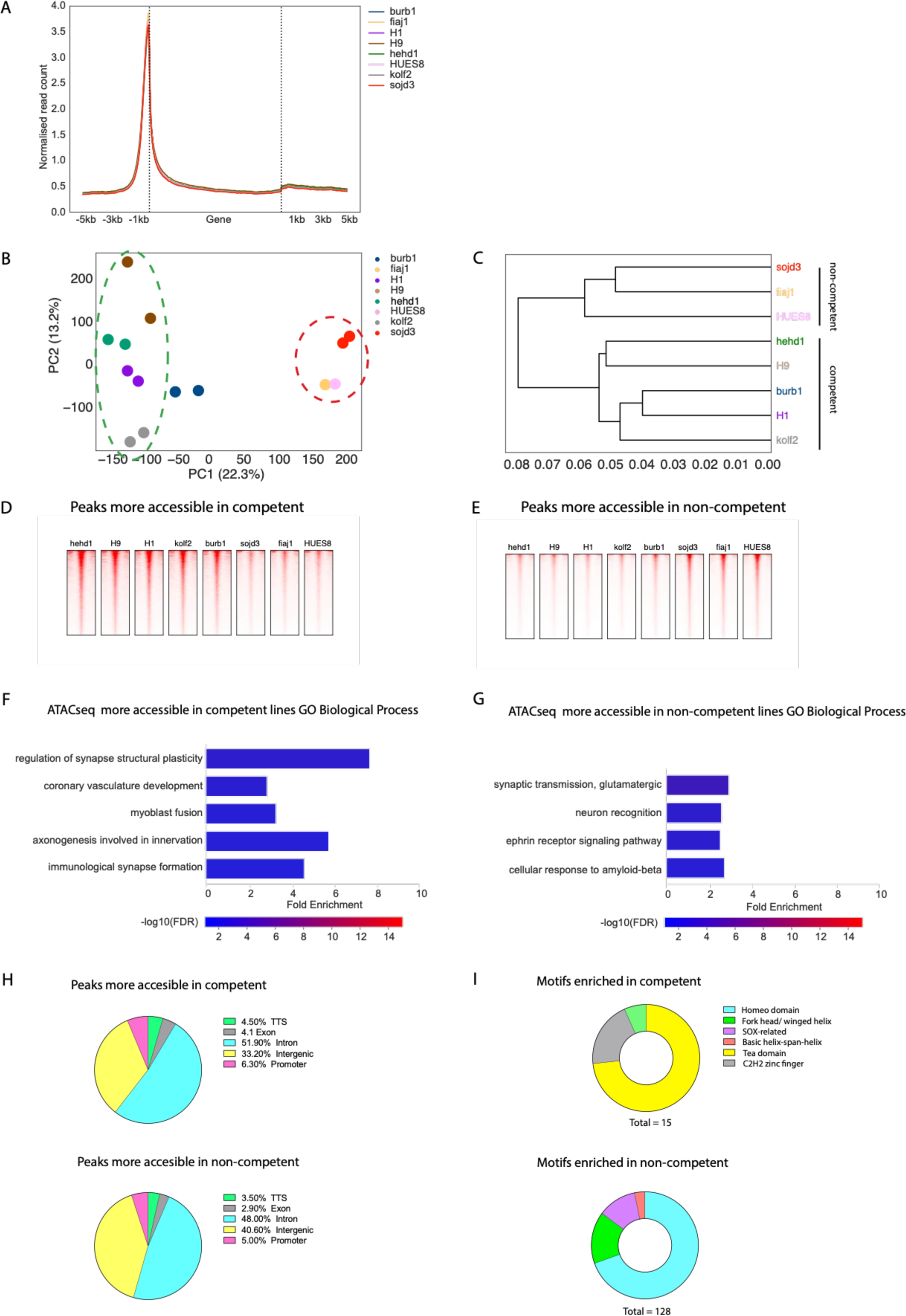
A - metagene plot of chromatin accessibility spanning a window between 5kb upstream from transcription start sites and 5kb downstream from transcription termination sites showing very similar profiles in all the cell lines tested, B – scatterplot of the two first principal components of the PCA of quantified differential ATAC-seq peaks in iPSC and ESC lines (220,897 variable peaks), encircled in green are competent lines and in red are the non-competent lines, C – hierarchical clustering plot of the quantified differential ATAC-seq peaks in iPSC and ESC lines (220,897 variable peaks), note that burb1 clusters with competent despite being functionally classed as a non-competent line, D – heatmaps of the ATAC-seq signal for peaks more accessible in competent cell lines separated by cell line, E - heatmaps of the ATAC-seq signal for peaks more accessible in non-competent cell lines separated by cell line, F - bar plot of selected significantly enriched terms from Panther GO BP analysis of genes within 2kb downstream of peaks differentially more accessible in competent lines, G - bar plot of selected significantly enriched terms from Panther GO BP analysis of genes within 2kb downstream of peak differentially more accessible in non-competent lines, H – circle charts summarizing genomic location of differentially accessible peaks more accessible in competent (top) or non-competent (bottom) lines, I – circle charts summarizing classification of transcription factors whose binding motifs were significantly (E value <10- 30) enriched in differentially accessible peaks more accessible in competent (top) or non-competent (bottom) lines.

To connect the changes in the chromatin accessibility with a biological function, we related differential peaks to the closest gene within 2kb downstream to identify more accessible promoters typically associated with active transcription.^46^ Surprisingly, GO term analysis did not reveal any clear patterns in the differentially accessible promoters, as both more accessible in competent and more accessible in non-competent groups contained terms associated broadly with multiple system development (Figures 2F-G, Table S3). We then analysed the distribution of differentially accessible peaks, which showed that the vast majority localised to intergenic regions and introns, rather than promoters, pointing to putative regulatory elements (REs) (Figure 2H).

We next performed motif enrichment analysis of differentially accessible peaks, which returned motifs specific to homeodomain, fork head (FOX) and SOX factors with important roles in organ development and tissue differentiation (Figure 2I, Table S4). The binding of transcription factors to REs in the genome is crucial for orchestrating the complex expression patterns of developmental genes.^47^ Changes in RE accessibility drive cell fate transitions, ultimately shaping a unique enhancer landscape that defines cellular identity. ^48,49^ Notably, Wnt signaling promotes the activation of enhancers specific to caudal development, while non-gastrulating tissues maintain an enhancer landscape akin to that of the epiblast.^50–53^ This distinction offers a plausible mechanism for WNT-mediated memory in non-competent lines and, corroborated by the differences in transcriptome and the effects of WNT inhibition, supports the hypothesis that non-competent lines may reflect a more posterior epiblast-like character even before pluripotency exit and onset of differentiation.

### Restoring uncommitted primed chromatin state

Since the differences between individual cell lines were retained at the chromatin level, we explored the possibility of restoring the chromatin landscape of non-competent lines to a competent state. Under natural circumstances chromatin remodelling associated with cell fate transitions is permanent and irreversible^54–56^. However, upon overexpression of individual or cocktails of transcription factors, exposure to undifferentiated cell cytoplasm or use of small molecules acting on chromatin modifiers, the cell chromatin and identity can be altered.^57,58^ The global chromatin landscape is reset during reprogramming of somatic cells to pluripotent stem cells,^59^ and also in the transition from primed to naïve pluripotent state.^60,61^ We reasoned that findings from the field of somatic cell reprogramming could inform strategies to restore a competent chromatin state in non-competent lines. From a practical standpoint, we aimed to establish a transgene-free approach that could be applied to a broad range of PSC lines. Such approaches have been successfully applied in resetting primed pluripotency to the naïve state and, more recently, in reprogramming somatic stem cells to pluripotent stem cells.^62,63^

We adapted part of a recently published stepwise method of chemical reprogramming of somatic cells to iPSCs.^63^ The last step (Step IV) of this method reprograms extraembryonic-like POU5F1+ epithelial precursors to chemically induced PSCs and contains molecules that are also used for primed to naïve resetting, making it suitable to apply on already pluripotent cells.^62^ We first applied the unaltered Step IV of the Guan et al. method to our pluripotent cells. This produced viable colonies with typical PSC morphology; however, their cerebral organoid competency was not improved, and the derived lines produced disorganised tissues without typical neural buds (Figure S2 A).

We therefore sought to develop a refined approach that is more specific to reversing the caudal epiblast chromatin landscape. Since the caudal epiblast is pre-determined to gastrulate, we analyzed the epigenetic changes underlying and potentially preceding gastrulation to determine which other aspects of chromatin we should target.^64–67^ Step IV of the Guan et al. protocol already involved molecules that modify epigenetic regulators of gastrulation but with the exception of H3K9 methylation.^67^ During gastrulation H3K9me3 is deposited in gene bodies, promoters and transcription termination sites, which in turn drive early endoderm and mesoderm germ layer specification.^68^ In general, during cell fate transitions, H3K9 methylation represses lineage-specific promoters and enhancers ^48,69,70^. Interestingly, aberrant H3K9 methylation can also arise as an artifact of *in vitro* cell culture due to exposure to high concentrations of growth factors, which is relevant for our experimental model.^69,71^ We therefore included a H3K9-methylase inhibitor and lowered the concentration of tranylcypromine, which could prevent removal of H3K9 methyl groups (Figure 3B).

**Figure 3.**
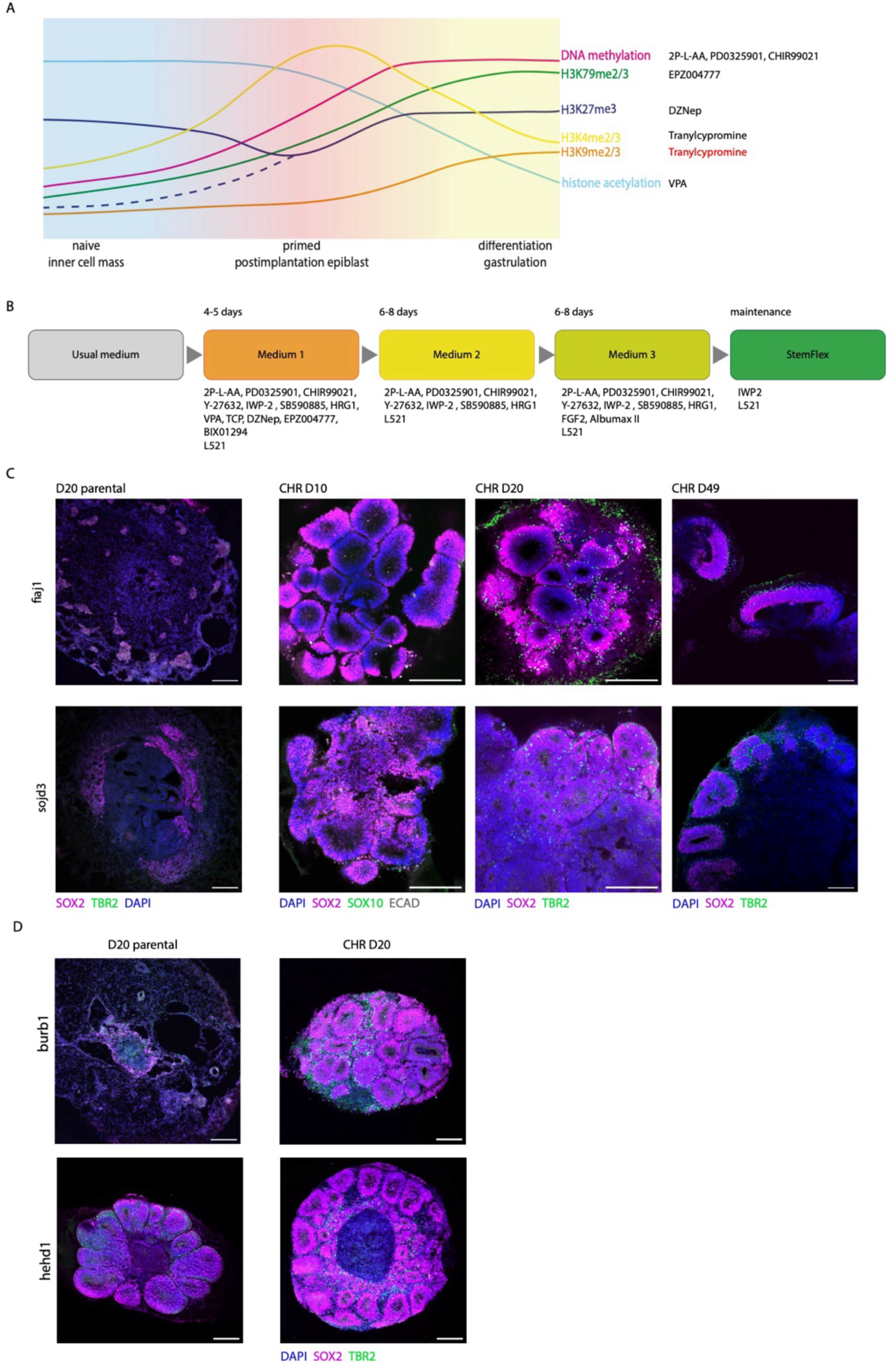
A - summary of epigenetic changes related to cell fate transition between blastocyst stage and gastrulation and their *in vitro* counterparts; the split in the purple line representing levels of H3K27me3 show the differences between *in vivo* (dashed line) and *in vitro* (solid line) data; on the right the names of small molecules from Step IV are listed next to the epigenetic change they target; the counterproductive action of tranylcypromine on H3K9me2/3 is highlighted in red, B – schematic of the chromatin restoration treatment with small molecules; key media components or specific culture conditions and typical duration listed for each step, C - representative images of cerebral organoids generated from non-competent lines fiaj1 and sojd3 before (leftmost image, organoid on day 20) and after CHR treatment (3 images on the right of organoids analysed on day 10, 20 and 49, respectively), showing morphology and marker expression improvement and correct developmental trajectory in time, scale bars 200 μm, D – representative images of day 20 organoids from parental (untreated, left) and CHR (treated, right) non-competent line burb1 and competent line hehd1, showing improvement in morphology and marker expression in burb1 after CHR and similar morphology and marker expression in hehd1 after CHR, scale bars 200 μm.

When we applied this chromatin restoration (CHR) method to two of the non-competent lines that were resistant to other attempts, sojd3 and fiaj1, we were able to recover colonies that could generate typical cerebral organoids (Figure 2S B). Organoids generated with this method showed the correct morphology, correct marker expression and correct developmental progression (Figure 3C).

We next tested whether this method is generally applicable to restore competency of other PSC lines. We applied the same treatment to the non-competent burb1 line, as well as hehd1 to assess any unwanted side effects of CHR on a naturally competent line. Organoids generated from colonies recovered after CHR treatment of burb1 and hehd1 lines were of high quality, confirming that CHR restored competency of burb1 to generate rostral neural structures, and that the competency of hehd1 cells was not compromised (Figure 3D).

We next asked whether CHR removed posterior bias and maintained general pluripotency or simply skewed differentiation potential towards anterior identities, by analyzing the pluripotency state of the CHR lines. To confirm pluripotency, we first looked at the expression of canonical pluripotency markers. All the CHR lines expressed the canonical pluripotency markers SOX2, OCT4 (POU5F1) and NANOG both at the mRNA level (Figure 2S C), and also at the protein level as detected by immunofluorescence (Figure 2S D).

To functionally confirm that the lines were able to differentiate into all three germ layers, we generated two more types of organoids: kidney - a mesoderm derivative, and intestine –an endoderm derivative. The hallmark of successful kidney organoid differentiation is epithelialisation of mesenchymal cells and generation of tubules, already visible on day 14 in phase contrast microscopy.^72^ We generated kidney organoids from all 4 CHR lines and their parental counterparts, and analysed resulting tissues for the presence of epithelial tubes. We stained representative samples at day 15 to confirm correct marker expression and location (Figure 2S E) and quantified the fraction of organoids with tubules in each parental and CHR cell line (Figure 2S F). As expected, all CHR lines produced kidney organoids with correct morphology and marker expression, whereas of the parental lines only hehd1 produced optimal tissues. In contrast, the non-competent parental lines burb1 and fiaj1 produced homogenous mesenchymal aggregates that did not show areas of correct morphology or marker expression, whereas sojd3-derived tissues tended to disintegrate before they could be analysed (Figure 2S E-F).

When differentiated into intestinal organoids, the CHR sojd3 and burb1 lines produced tissues with widespread areas positive for epithelial E-cadherin and Villin-1, with some scattered SOX17 staining (Figure 2S G). Co-staining for E-cadherin and CDX2 revealed a large proportion of double-positive tissues, a hallmark of hindgut (Figure 2S H).^73^ The parental lines on the other hand generated more compact tissues largely negative for E-cadherin and lacking specific Villin-1 staining, although some areas in organoids from parental burb1 lines showed correct marker expression. Thus, we confirmed that the CHR protocol we established here completely restored pluripotency and full differentiation potential towards all three germ layers.

### CHR treatment restores the anterior uncommitted epiblast state

Since the CHR method was able to restore cellular competency to produce anterior neural tissues, we next investigated whether the previously observed differences in chromatin accessibility and gene expression were reversed. We first analysed the chromatin accessibility pattern of CHR lines and their parental counterparts with ATAC-seq in the burb1, hehd1, fiaj1 and sojd3 lines. As expected, PCA revealed that the CHR lines clustered closer together with the naturally competent line hehd1 when 225,756 variable peaks were considered (Figure 4A and Figure 3SA). We then examined how regions differentially accessible between competent and non-competent lines behaved after chromatin restoration (Figure 4B-C). The accessibility of the peaks in CHR lines correlated with the accessibility in competent lines – peaks more accessible in competent lines were also more accessible after CHR (Figure 4B), whereas peaks less accessible in competent lines became less accessible with CHR (Figure 4C). Interestingly, the changes in burb1 were less profound than in fiaj1 and sojd3, in line with the fact that before treatment burb1 chromatin accessibility differed the least from competent lines (Figure 2B). The accessibility of chromatin in a naturally competent line hehd1 changed the least and when peaks differentially accessible between competent and non-competent cell lines were compared, the difference was minimal. This shows that for chromatin accessibility, the endpoint of CHR is similar irrespective of the chromatin landscape of the starting cell line, and in line with the functional outcome.

**Figure 4.**
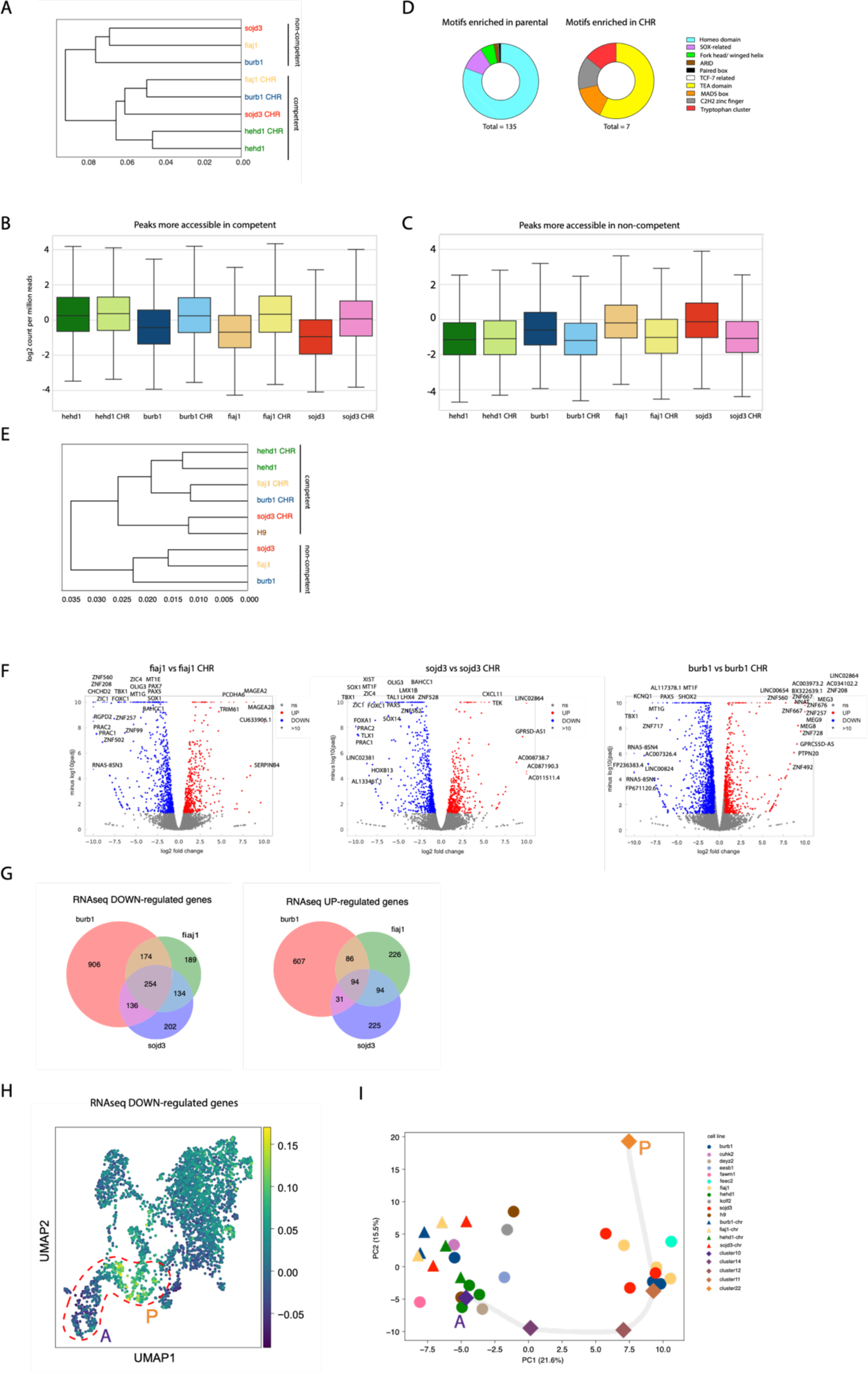
A - hierarchical clustering plot of the quantified ATAC-seq peaks in iPSC lines before (parental) and after treatment (CHR) showing higher similarity between competent line hehd1 and all CHR lines than between CHR lines and their parental counterparts (analysis 225,756 top variable peaks), B - box plot summarizing changes in chromatin accessibility in parental and CHR lines in regions previously defined as peaks more accessible in competent lines, C - box plot summarizing changes in chromatin accessibility in parental and CHR lines in regions previously defined as peaks more accessible in non-competent lines, D - circle charts summarizing classification of transcription factors whose binding motifs were significantly (E value<10-30) enriched in differentially accessible peaks more accessible in parental (left) or CHR (right) lines, E - hierarchical clustering plot of the bulk transcriptome of parental and CHR lines (differential genes defined by VST threshold), F - volcano plots of differentially expressed genes between CHR and parental cell lines for fiaj1 (left), sojd3 (middle) and burb1 (right), G – overlap of differentially expressed genes in CHR vs parental between the fiaj1, burb1 and sojd3 cell lines, left – down regulated genes, right – up regulated genes, H – day 8 SEM feature plot of combined 254 transcripts consistently downregulated after CHR in burb1, fiaj1 and sojd3, showing that these transcripts are more abundant in the posterior epiblast, encircled is the region of the UMAP corresponding to the epiblast clusters, I - scatterplot of the first two principal components of the PCA of the bulk transcriptomes of H9 ESC line, competent and non-competent iPSC lines, parental and CHR lines and “pseudo-bulked” epiblast clusters of the day8 SEM, (analysis of 250 top variable genes).

Analysis of the transcription factor binding motifs showed that the peaks that closed in restored lines were mainly enriched in the same class of motifs as those found in peaks more accessible in non-competent lines; predominantly the homeodomain and SOX-related transcription factors (Figure 4D, Table S5). Collectively, this suggests that the chromatin restoration reversed the chromatin accessibility changes that distinguish competent and non-competent lines.

We then investigated whether CHR treatment also corrected the aberrant gene expression in non-competent cells. We performed bulk RNA-seq on all CHR lines and their parental counterparts, and included a naturally competent ES line, H9, for comparison. Principal component analysis of the transcriptomic data showed that CHR lines clustered close together with naturally competent lines hehd1 and H9 (Figure 4E, Figure 3S B).

To define genes whose expression was affected by the CHR treatment, we performed DE-seq analysis of genes that changed in expression in burb1, fiaj1 and sojd3 upon CHR treatment (Figure 4F, Table S6). Several hundred genes were up- or down-regulated in each of the non-competent lines after CHR treatment. We next identified the overlapping downregulated (core DE down) and upregulated genes (core DE up) shared between the burb1, fiaj1 and sojd3 lines (Figure 4G). Comparison of DE genes between CHR and parental lines and competent and non-competent lines showed a high degree of consistency between transcript levels in CHR and competent lines (Figure 3S C, Figure 1C). When we interrogated the bulk RNA-seq dataset of competent vs non-competent cell lines (Figure 1B), we found that the core DE down genes were also highly expressed in another non-competent line, feec2 (Figure 3S D).

Finally, we investigated whether the CHR process aligns with the epigenetic regionalisation that we postulated was ongoing in the non-competent lines. We returned to the SEM reference dataset to interrogate which clusters expressed the genes that changed with CHR. Strikingly, core DE down genes were overrepresented in the posterior clusters 11,12 and 22 (Figure 4H). We then generated pseudo-bulked objects of the relevant clusters of the SEM scRNA-seq dataset to perform PCA analysis together with all RNA-seq datasets used in this study so far (competent, non-competent and CHR lines). Dimensionality reduction analysis again showed that PC1 captured the anterior to posterior gradient of the SEM epiblast and revealed separation of competent, CHR and non-competent lines alongside the same axis (Figure 4I). The anterior-most cluster 10 localised the closest to the competent and CHR lines, anterior cluster 14 occupied an intermediate territory, whereas posterior clusters 11,12 and 22 segregated with non-competent lines. Collectively this indicates that the transcriptomic differences between non-competent vs competent and parental vs CHR lines both capture the developmental oscillation around the antero-posterior epiblast regionalisation point.

### CHR treatment restores bivalent histone marks

ATAC-seq analysis showed that the areas of differentially accessible chromatin between competent and non-competent lines localised to intergenic and intronic regions, consistent with regulatory elements, with only a minor fraction localised in promoters (Figure 2H). However, having identified the core genes whose expression was responsible for the caudalised phenotype, we investigated if promoter chromatin accessibility could explain the differences in their expression. We analysed chromatin accessibility in the promoters of core DE genes after CHR in burb1, fiaj1 and sojd3 lines (Figure 4G). To our surprise, we did not find a correlation between chromatin accessibility and transcript levels of differentially expressed genes, with only a few genes downregulated after CHR showing a more closed conformation (Figure 5A). Interestingly, when we looked at the individual ATAC-seq and RNA-seq tracks of the core DE genes, we noticed that although the RNA expression was markedly decreased after CHR, chromatin accessibility did not change substantially, and the promoter regions of DE genes still showed accessible conformation irrespective of transcriptional status (Figure 4S A). This was consistent with the chromatin differences that determine competency mainly influencing distal RE but could not explain how the differences in expression of those genes are mediated at the chromatin level.

**Figure 5.**
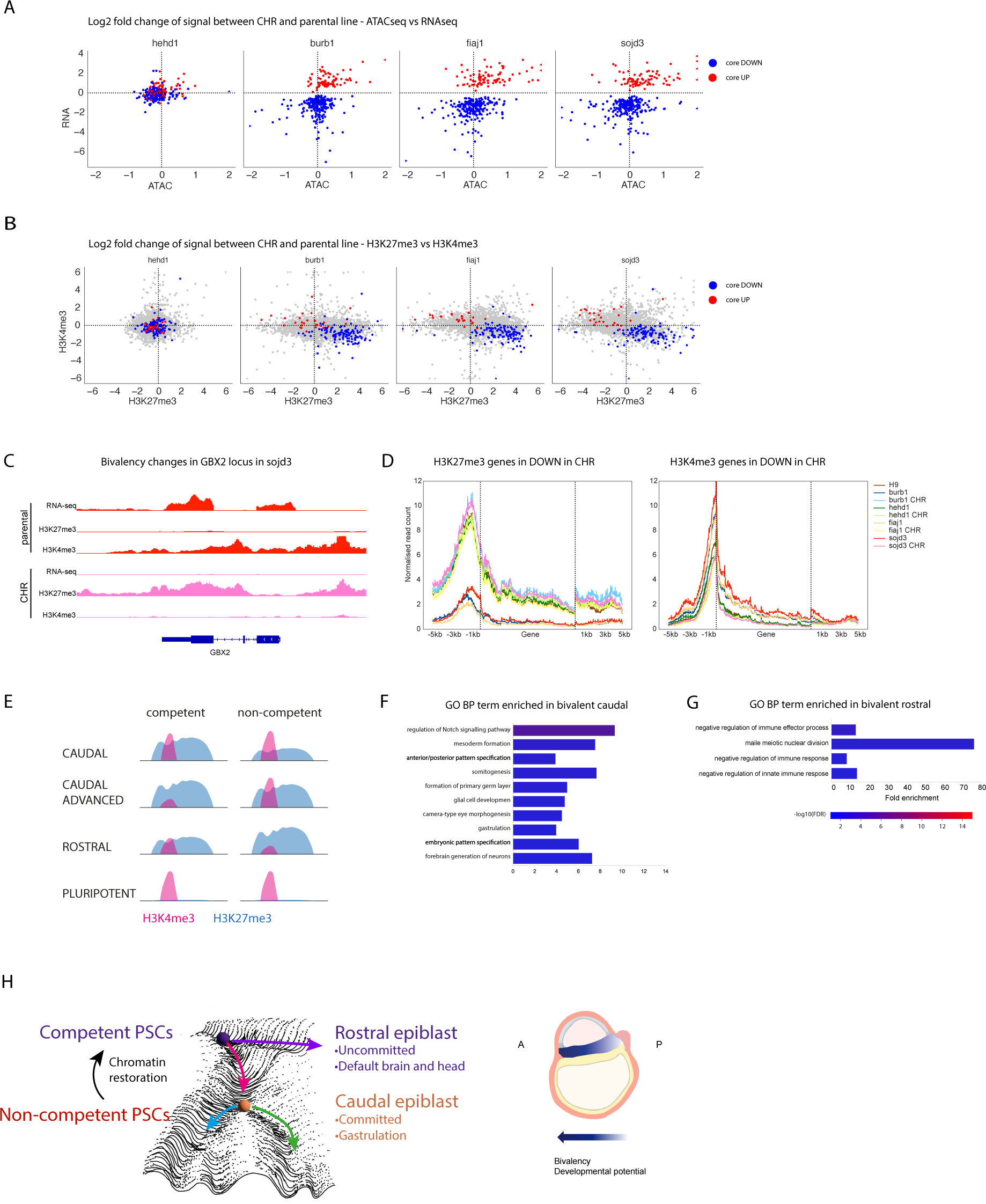
A -scatterplots of log2 fold change of ATAC-seq signal between CHR vs parental lines vs log2 fold change of RNA-seq signal between CHR vs parental lines, blue datapoints represent overlapping downregulated transcripts in CHR burb1, fiaj1 and sojd3 lines (core DE DOWN), red datapoints represent overlapping upregulated transcripts in CHR burb1, fiaj1 and sojd3 lines (core DE UP), B-scatterplots of log2 fold change of H3K27me3 CUT&Tag signal between CHR vs parental lines vs log2 fold change of H3K4me3 CUT&Tag signal between CHR vs parental lines; blue datapoints represent overlapping downregulated transcripts in CHR burb1, fiaj1 and sojd3 lines (core DE DOWN), red datapoints represent overlapping upregulated transcripts in CHR burb1, fiaj1 and sojd3 lines (core DE UP), grey datapoints represent all other bivalent promoters, C – Integrative Genomics Viewer snapshot of RNA-seq, H3K4me3 and H3K27me3 tracks of the GBX2 locus in sojd3 and sojd3 CHR lines, D – metagene profiles of H3K27me3 (left panel) and H3K4me3 (right panel) in parental and CHR iPSC lines and H9 ESC line, of the core DE DOWN genes, E – expected patterns of H3K27me3 and H3K4me3 in promoters of genes involved in rostro-caudal pattering and pluripotency, E - bar plot of selected significantly enriched terms from Panther GO BP analysis of genes classified as caudal based on the H3K27me3 and H3K4me3 balance in the 2kb upstream promoter region, F - bar plot of selected significantly enriched terms from Panther GO BP analysis of genes classified as rostral based on the H3K27me3 and H3K4me3 balance in the 2kb upstream promoter region, H-graphical summary of the findings in this study integrating caudalisation in the non-competent lines and chromatin restoration treatment

Many developmental genes with low DNA methylation and high chromatin accessibility show a bivalent signature of histone H3 lysine 27 trimethylation (H3K27me3) repressive and histone H3 lysine 4 trimethylation (H3K4me3) activating chromatin marks in their promoters that enables quick transcriptional activation upon initiation of differentiation programmes.^74^ We therefore analysed H3K27me3 and H3K4me3 modifications by CUT&Tag^75^ in parental and CHR lines. We first correlated core UP and DOWN regulated gene mRNA expression with intensity of signal for H3K27me3 and H3K4me3 in their promoter regions (Figure 5B). In the CHR lines the core DE DOWN genes (blue datapoints) were enriched in H3K27me3 and depleted in H3K4me3 marks in comparison to their parental counterparts. On the other hand, core DE UP genes (red datapoints) showed H3K27me3 depletion and H3K4me3 enrichment. This effect was specific to the DE genes, since there was no global trend of enrichment or depletion of these chromatin marks in all expressed genes (grey datapoints). As expected, core DE genes did not show a shift of H3K4me3 or H3K27me3 in the naturally competent line hehd1 after CHR treatment.

We next visualised chromatin mark changes at individual gene loci. The caudal patterning gene GBX2 exhibited decreased H3K27me3 and elevated H3K4me3 in the sojd3 parental cell line that was reversed upon CHR treatment (Figure 5C). Global analysis of H3K27me3 signal over the core downregulated genes in parental non-competent lines confirmed its depletion, particularly in the promoter regions (Figure 5D). In contrast, CHR lines displayed H3K27me3 profiles more similar to naturally competent hehd1 and H9 lines. When the whole genome was analysed, parental fiaj1 and burb1 lines showed slightly lower global level of H3K27me3 in all the promoters than competent and CHR lines, whereas the difference was minimal in the parental sojd3 line (Figure 4S B-C). H3K4me3 was enriched in the core DE DOWN genes in non-competent lines (Figure 5D) but not globally (Figure 4S B-C). The enrichment of H3K4me3 in promoter regions in core DE DOWN was the strongest in the sojd3 line (Figure 5D). Collectively this pointed to the expression of DE genes being regulated by the balance of H3K27me3 repressive marks and H3K4me3 activating marks, where low H3K27me3 and high H3K4me3 status predicted expression and competency.

Interestingly, we did not observe pronounced differences in DNA methylation in CHR vs parental lines nor in naturally competent vs non-competent lines, consistent with the notion that the differentially regulated regions were already hypomethylated and that the main regulatory mechanism works at the level of histone post-translational modifications (Figure 4S D-H). When we focused on the genes downregulated in restored lines, we did not detect a clear trend of increased or decreased DNA methylation that would correlate with gene expression (Figure 4S F). Although some DNA regions became hypomethylated in the CHR cells, they were not consistent between cell lines and only 15 hypomethylated regions overlapped with core DE down genes (Figure 4S G-H). Overall, we conclude that the loss of chromatin bivalency in a subset of developmental genes in the non-competent lines results in their aberrant expression that underpins the restricted potential in these lines. The CHR treatment restores developmental capacity by counteracting bivalent chromatin erosion and resetting the transcription levels of key developmental regulator genes.

### Bivalency patterns reveal epiblast developmental asynchrony

The balance of active H3K4me3 and repressive H3K27me3 chromatin marks emerged as the most immediate, sensitive and dynamic readout of the cell state. We therefore hypothesized that by analysing the pattern of H3K4me3 and H3k27me3 directly, we would be able to reveal the earliest embryonic gene set that drives epiblast A-P axis establishment.

To define the pool of genes involved in this fate transition we mapped genes whose promoters show bivalency in at least one of the lines (all possible bivalent). We then speculated that genes showing a decrease in H3K27me3 in the fiaj1, burb1 and sojd3 non-competent lines would be the caudal specific genes, whereas the genes showing a gain in levels of H3K27me3 would be rostral specific genes (Figure 5E). Finally, genes bivalent in H9 and hehd1 but more active (decreased H3K27me3and increased H3K4me3) in non-competent lines would be the genes that drive caudalisation. We performed DE-seq analysis on all possible bivalent genes using levels of histone modifications in H9 cell line as a reference (Table S8). As a control we checked the chromatin status of pluripotency genes to confirm they showed strong signal for the active chromatin mark H3K4me3 in all the lines (Figure 5S A).

We classified the genes as belonging to one of the groups defined above and performed GO term analysis for each of the groups. As expected, we found that the predicted caudal genes (loss of H3K27me3 in non-competent) were enriched in BP terms associated with gastrulation and mesendoderm development (Figure 5F, Table S9). The genes where H3K27me3 loss was accompanied by H3K4me3 gain partially overlapped with core DE down in CHR lines (Figure 5S B). Interestingly, several of those genes showed an unexpected expression pattern in the developing embryo (Figure 5S C-E). They were highly expressed in pre-implantation epiblast, lower in CS7 epiblast and re-emerging in primitive streak and developing mesoderm, suggesting their downregulation in primed epiblast.^34^ Importantly, we detected a number of genes that showed higher H3K27me3 signal in the non-competent parental lines, supporting a specific mechanism consistent with regionalisation rather than general reduction of H3K27 trimethylation (Table S8). We hypothesized that loci higher in H3K27me3 in non-competent would include genes associated with anterior neural development; however, this group was surprisingly enriched in genes involved in meiosis (Figure 5F). Those genes are relevant to germ line development, which in mouse is possible in the short window of formative pluripotency.^20,76^ However, our CHR cell lines did not show higher expression of formative pluripotency markers than parental lines, consistent with the typical primed conditions they were cultured in and with the fact that human primed PSCs are germ layer competent.^77^ When naturally competent lines were compared to non-competent lines, the competent lines tended to express higher levels of primed markers (Figure 5S H-I).^78^ This suggests that anterior competent cells occupy a state of early primed pluripotency, which is competent to produce anterior brain structures and might have enhanced potential for germ line, and that the pluripotency spectrum exists both in spatial and temporal dimensions.

## Discussion

Human primed pluripotent stem cells capture the unique state of the developing embryo when the epiblast is competent to respond to a complex network of patterning signals to establish a huge repertoire of different cell fates needed to build a body. Our study uncovers how initial epiblast regionalisation is encoded at the epigenetic level and is compatible with pluripotency maintenance. We experimentally establish that the genes involved in possible fate choices have high chromatin accessibility and are receptive to regulation by external signalling. These genes respond by balancing active and inactive chromatin marks around the point of bivalency to finally resolve the chromatin state in one of the directions. The correct pattern of bivalency is recognised as being crucial for pluripotency, and bivalency on key developmental genes is a predictor of differentiation capacity.^20,52,79,80^ The balance of repressive and active chromatin marks therefore emerges as the most accurate predictor of gene activity, surpassing chromatin accessibility and DNA methylation status,^81–85^ whereas unguided cerebral organoid differentiation competency transpires as a readout of genuine pluripotency.

Interestingly, developmental genes that are targets for the PRC2 complex, the molecular machinery responsible for the repressive mark of bivalency H3K27me3, are the most variably expressed amongst PSC lines, often varying in different clones from genetically identical donors and showing little correlation with the genetic background.^86^ This is in line with our conclusions that developmental genes are particularly sensitive to transient patterning signalling, which persistently changes proportions of H3K27me3 and H3K4me3 marks, thus inducing lineage priming.

Since the lineage primed states are persistently encoded at the epigenetic level but do not dissolve the pluripotency network, they could explain the mechanism of non-genetic differentiation bias in cell lines that otherwise fulfil pluripotency criteria. This would result from a physiological mechanism of epiblast regionalisation that is not fully controlled in the *in vitro* environment and not subjected to safeguarding mechanisms present in a complete embryo. In this study, we demonstrate that the differentiation bias deposited in the form of H3K27me3 and H3K4me3 imbalance can be reversed by the treatment with small molecules and maintained in the most competent (highest unbiased developmental potential) state by maintenance of pluripotent stem cell culture in conditions that minimise the strongest epiblast patterning signals, namely WNT and BMP.^31^

The footprint of variable H3K27me3 reflects the past signalling pressure and provides a snapshot of the patterning milieu even before fate transition occurs. An analogous mechanism was observed in neural progenitor cells (NPCs), where epigenetic memory is formed through antagonism between PRC2 and activating signalling inputs.^87^ Even transient expression of an active transcription factor in NPCs was able to overcome PRC2 repression and leave an epigenetic footprint that lasted over many cell divisions. Here, we demonstrate that a similar phenomenon in primed PSCs could be leveraged as a tool to probe the earliest effectors of differentiation by investigating a shift in H3K27me3 and H3K4me3 marks in promoters of the affected genes. This strategy allows the use of differential levels of bivalent marks to specify genes involved in early epiblast regionalisation. We also show that anterior competency involves retention of earlier developmental features, including bivalency on meiosis genes, consistent with the fact that specification of primordial germ cells in human and primate embryos precedes gastrulation.^88,89^

Overall, we paint a picture of asynchronous development of the human epiblast, where the anterior portion retains developmentally earlier features, whereas the posterior part undergoes priming for gastrulation (Figure 5H). The earliest decision is therefore a choice between gastrulation or maintenance of unchanged epiblast identity, with the latter being the default in the absence of signalling and permissive to anterior neural or brain fate. Interestingly, this initial asynchrony could be required to eventually align neural induction in the anterior and posterior epiblast to produce a continuous neural plate, whose induction through the neuromesodermal progenitor intermediate in the caudal embryo takes longer.^90^

A delay in fate transitions and maintenance of a developmentally early proliferative state has been described as a potential mechanism of expansion of the human forebrain^28^ but an analogous delay could start even earlier in embryonic development. In this context the huge reliance of anterior neural fate on PRC2-maintained bivalency might be a trade-off of the evolutionary expansion of the anterior nervous system in higher vertebrates. The stable primed epiblast that is needed to grow a large forebrain would need to extend the time window of bivalency on key genes and resist posteriorizing signals. This is supported by the observation that when PRC2 is supressed, naïve cells aberrantly upregulate mesoderm lineage upon priming and neural differentiation^67^, whereas primed pluripotent stem cells cannot acquire neural fate when PRC2 is supressed.^91–93^ The resulting uncoupling of anterior and posterior neural induction through this early epigenetic patterning could enable independent evolution of the anterior brain without affecting the posterior hindbrain or spinal cord.

## Acknowledgements

The Lancaster Lab is supported by the Medical Research Council (MC_UP_1201/9). SS was supported by a UKRI MRC Rutherford Fund Fellowship (MR/T016787/1), the Babraham Institute’s Epigenetics Strategic Programme Grant (BBS/E/B/000C0421) and a Career Progression Fellowship from the Babraham Institute. EW was supported by a BBSRC Industrial CASE training grant (BB/X511584/1).

We would like to thank MRC LMB Light Microscopy Facility and Scientific Computing. We would also like to thank Prof Austin Smith, Dr Ge Guo and Prof Ufuk Günesdogan for helpful discussions.

## Competing interests

MAL and MAS are co-inventors on a patent application entitled “Standardising pluripotent stem cells” (WO 2024/256607 A1) based on the method described in this paper. The patent application was filed by United Kingdom Research and Innovation. SS is a co-founder, shareholder and employee of Enhanc3D Genomics. MAL is an inventor on patents covering cerebral organoids and is co-founder and advisory board member of a:head bio.

## Supplementary figures

**Figure 1S.**
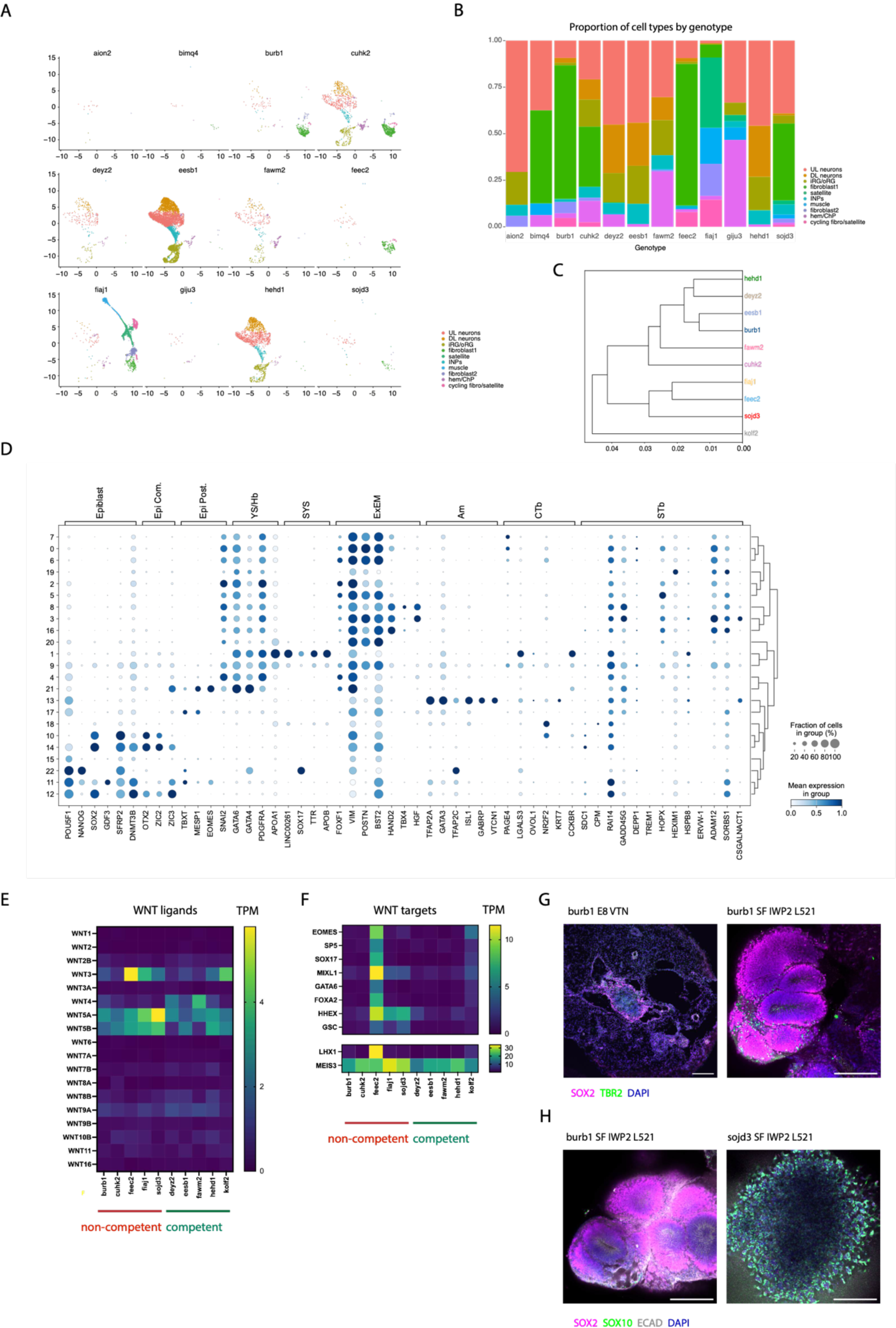
A - UMAPs of the pooled iPSC-derived cerebral organoids summarizing the organoid cell types split by donor genotype, B – stacked column plot of proportion of the cells types for each donor genotype, C - hierarchical clustering plot of the bulk transcriptome of competent and non-competent iPSC lines, D - dot plot illustrating the expression of key markers across the 22 cell type clusters, E – heatmap of normalised expression (transcripts per million (TPM)) of WNT ligands in competent and non-competent iPSC lines, F - heatmap of normalised expression (TPM) of canonical WNT targets in human PSCs in competent and non-competent iPSC lines, G – representative images of day 20 cerebral organoids generated from parental burb1 line and from burb1 cells cultured long term in Stemflex with IWP2, scale bars 200 μm, H – representative images of day 20 burb1 and day 10 sojd3 cerebral organoids generated from cells cultured long term in Stemflex with IWP2, scale bars 200 μm

**Figure 2S.**
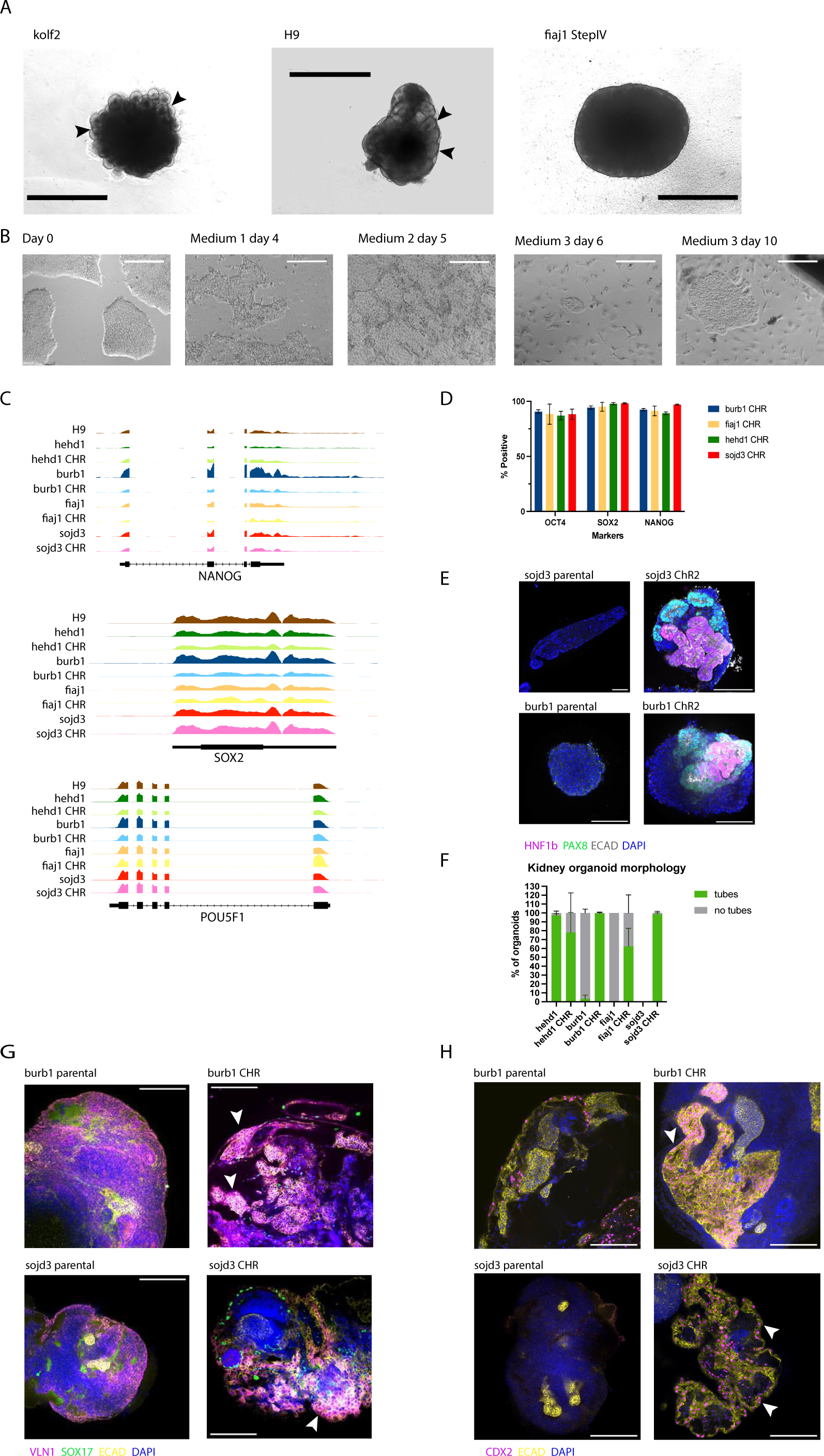
A – representative brightfield images of day 14 cerebral organoids generated from competent lines h9 or kolf2 or from fiaj1 cells recovered after Step IV of Guan et.al protocol^63^, scale bars 1mm, typical morphology with neuroepithelial buds (black arrowheads) visible in competent lines kolf2 and H9 but not in fiaj1, B - brightfield images of a culture undergoing CHR treatment, labelled above images with Medium stage and day since start of the treatment, C – Integrative Genomic Browser snapshots of RNA-seq signal in pluripotency markers loci (NANOG, POU5F1 and SOX2) in parental and CHR lines, and H9, D – column plot of proportion of cells positive for pluripotency markers by immunofluorescence (NANOG, POU5F1 and SOX2) in parental and CHR iPSC cultures, E-representative images of kidney organoids on day 15 generated from sojd3 parental and CHR lines and burb1 parental and CHR lines, scale bars 100 μm; please note that the aggregate captured in sojd3 parental kidney organoid preparation is an example of a rare event and that the DAPI-positive nuclei in this aggregate are smaller and more irregular than in other preparation suggesting necrosis;, F – quantification of bright field image morphology of day 13-15 kidney organoids from two batches, tissues with at least one area showing epithelial tube development were classed as “tubes”; please note that due to only a handful aggregates forming in batch 1 of sojd3 parental line and no aggregates present in the second batch, this experimental variant could not be quantified, G – immunofluorescent staining images of intestinal organoids generated from sojd3 parental and CHR lines and burb1 parental and CHR lines, white arrowheads indicate areas of VLN1 and ECAD staining overlaps, scale bars 200 μm, H – immunofluorescent staining images of intestinal organoids generated from sojd3 parental and CHR lines and burb1 parental and CHR lines, white arrowheads indicate areas of CDX2 and ECAD staining overlaps, scale bars 200 μm

**Figure 3S.**
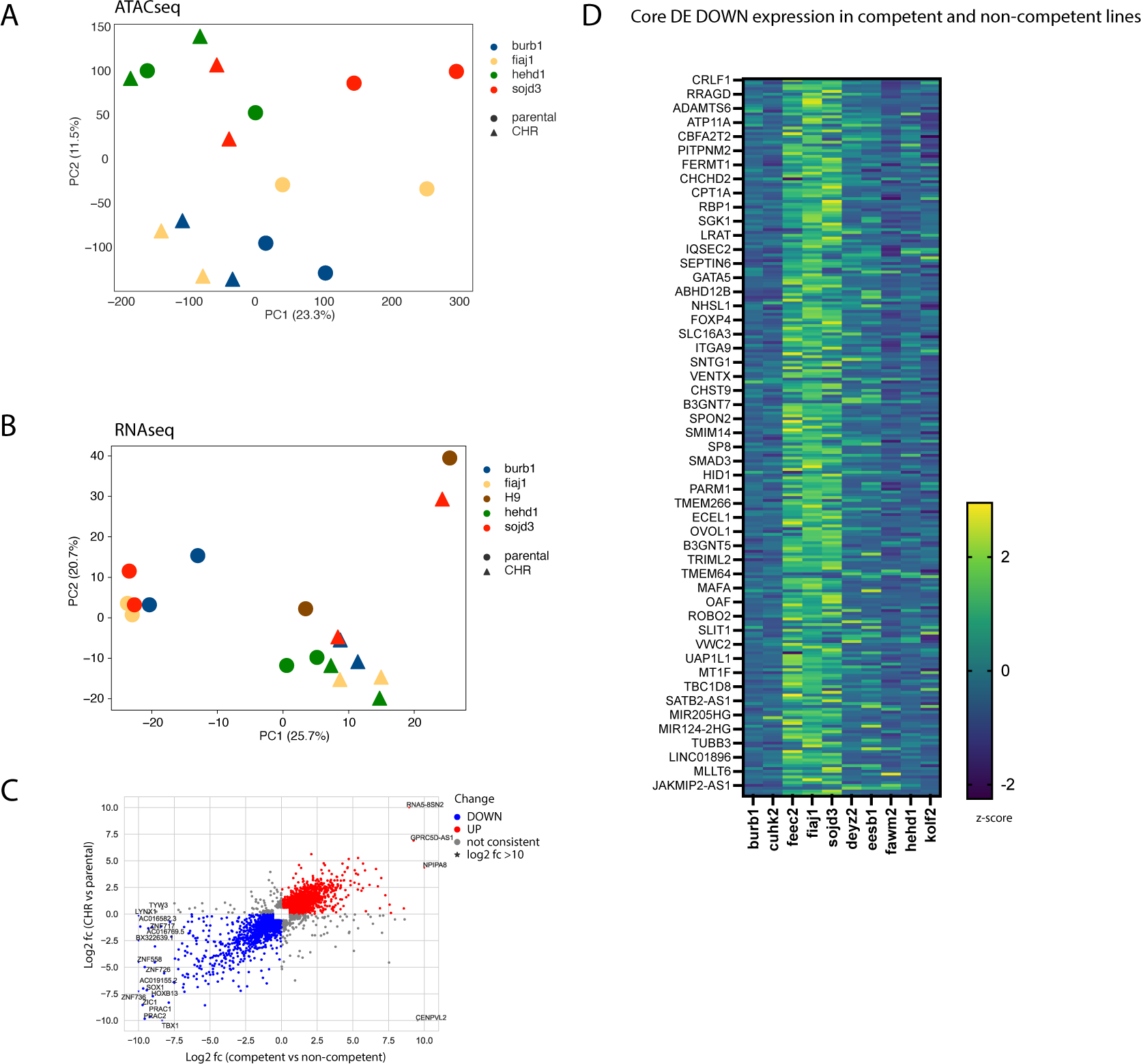
A - scatterplot of the two first principal components of the PCA of quantified ATAC-seq peaks in parental (circles) and CHR lines (triangles), based on 225,756 differential peaks, B - scatterplot of the two first principal components of the PCA of quantified bulk RNA-seq data in parental (circles) and CHR lines (triangles), differential transcripts based on VST threshold, C – scatterplot of log2 fold change of expression genes of genes identified as differentially expressed between competent and non-competent lines vs the log2 fold change of expression of those genes in CHR vs parental genes, blue datapoints highlight genes downregulated both in CHR vs parental and in competent vs non-competent, and red datapoints highlight genes upregulated both in CHR vs parental and in competent vs non-competent, grey datapoints highlight genes that do not show consistent change throughout both comparisons, D – heatmap of z-scored gene expression of 254 core DE DOWN genes (CHR vs parental) in the competent and non-competent iPSC lines used in this study

**Figure 4S.**
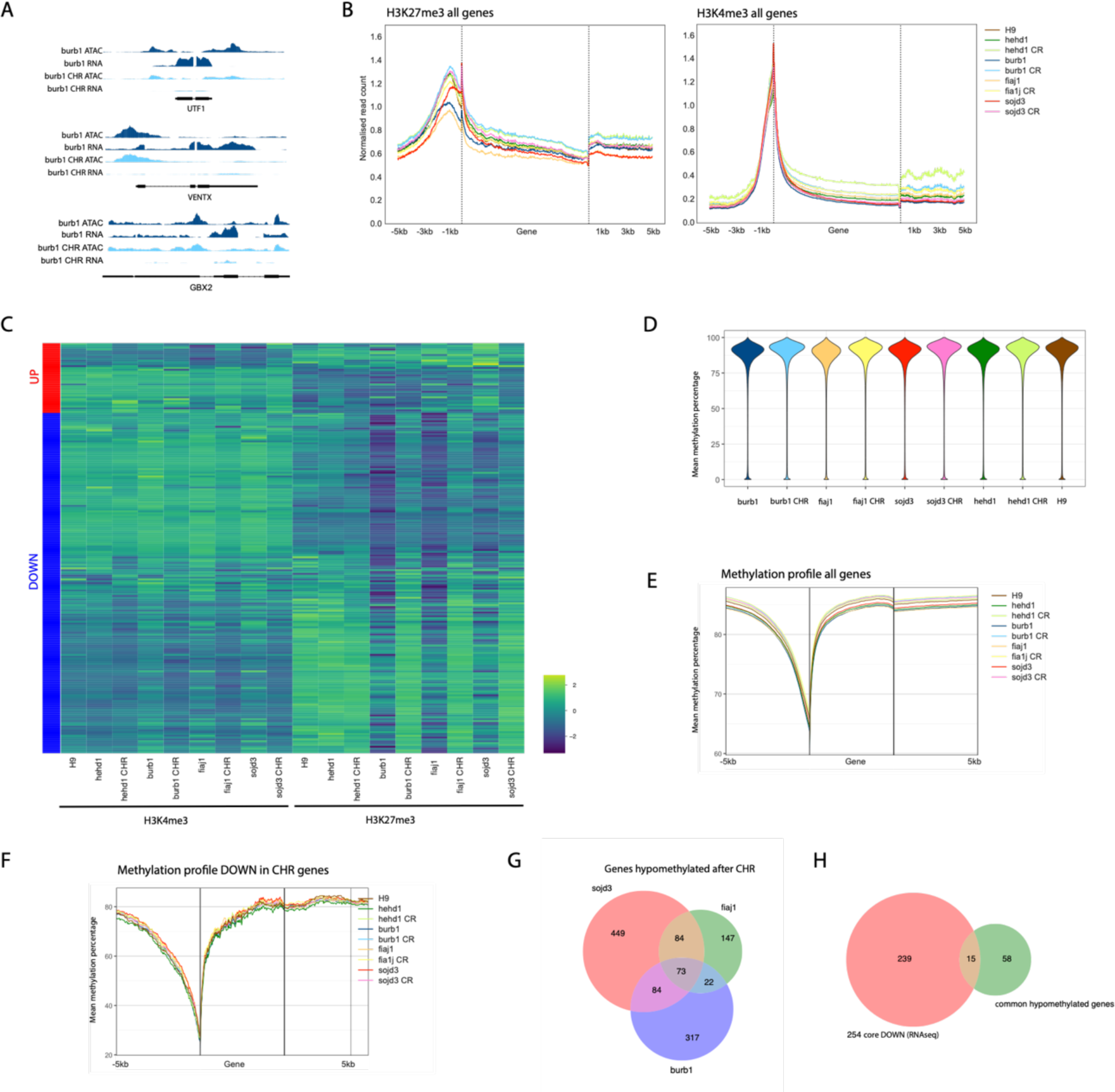
A - Integrative Genomics Viewer snapshot of ATAC-seq and RNA-seq tracks of the UTF1, VENTX1 and GBX2 loci in burb1 (parental) and burb1 CHR lines, B - metagene profiles of H3K27me3 (left panel) and H3K4me3 (right panel) in parental and CHR iPSC lines, and H9 ESC line, of all the bivalent genes spanning a genomic window between 5kb upstream from transcription start sites and 5kb downstream from transcription termination sites, C-heatmap of normalised signal values for H3K4me3 (left part) and H3K27me3 (right part) for promoter regions of the core DE UP and DOWN genes, D – beanplot of mean DNA methylation percentage in all loci genome wide, E-metagene profile of DNA methylation signal spanning a window between 5kb upstream from transcription start sites and 5kb downstream from transcription termination sites of all genes, F – DNA methylation metagene profile spanning a window of between 5kb upstream from transcription start sites and 5kb downstream from transcription termination sites of the core DE DOWN genes, G – overlap of genes hypomethylated after CHR between the three CHR lines burb1, fiaj1, and sojd3 to define common hypomethylated genes; genes were classed as hypomethylated if they overlapped with at least one differentially methylated region defined by at least 25% normalised methylation decrease after CHR, H-overlap of hypomethylated genes as defined in Figure 4S G with core DE DOWN

**Figure 5S.**
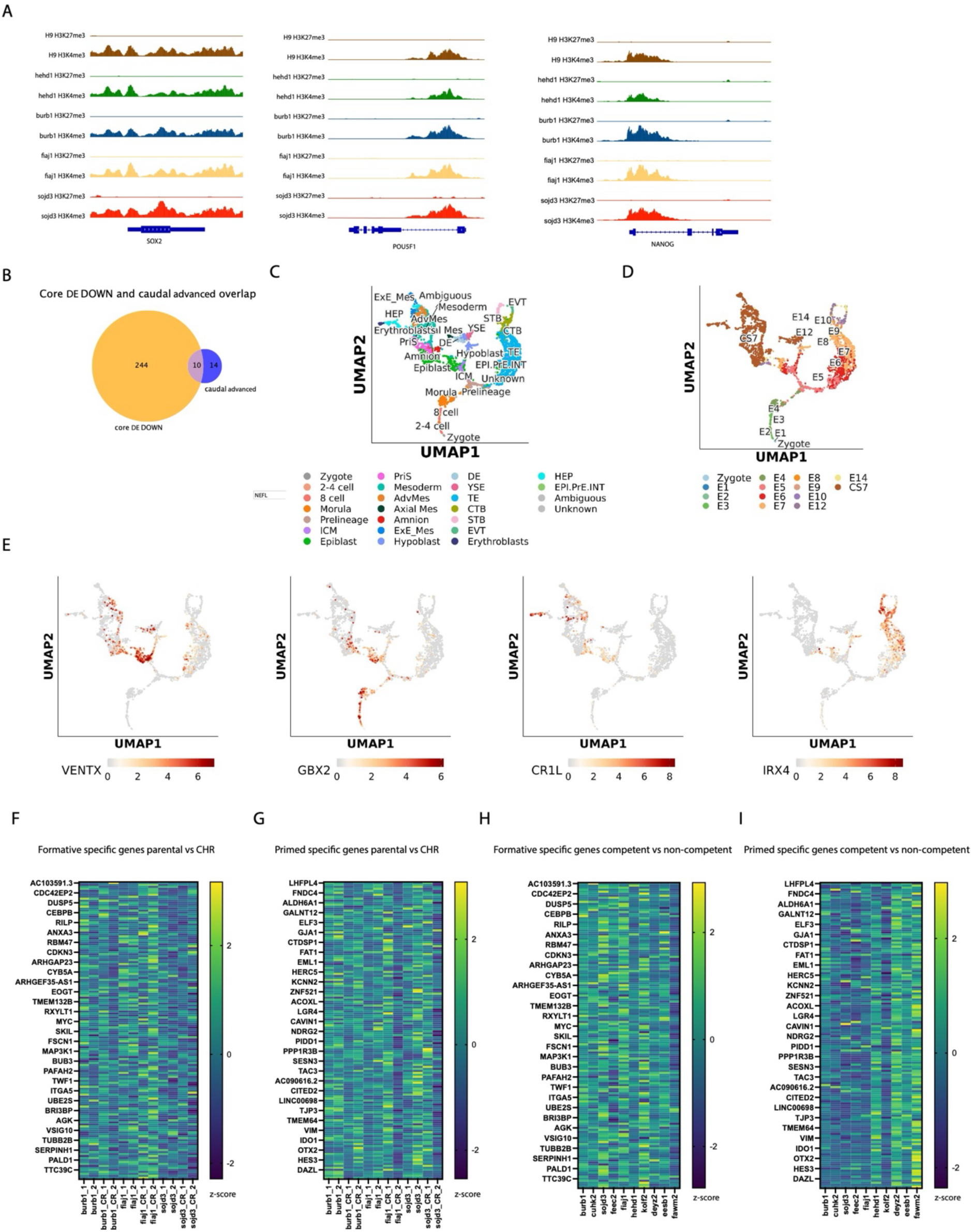
A - Integrative Genomics Viewer snapshot of H3K27me3 (top in each pair) and H3K4me3 (bottom in each pair) signals of all parental iPSC lines and H9 ESC line in the genomic loci of canonical pluripotency markers (SOX2, POUF5F1 and NANOG), showing high levels of active chromatin mark H3K4me3 and almost undetectable levels of H3K27me3 repressive mark, B – overlap of core DE DOWN genes and genes identified as “caudal advanced” through analysis of H3K27me3 H3K4me3 signal levels in bivalent promoters between H9 and non-competent lines, C – UMAP projection of the integrated human embryo reference datasets^34^ (generated using https://petropoulos-lanner-labs.clintec.ki.se/shinys/app/ShinyEmbryoRef) summarizing the cell identity, D – UMAP projection of the integrated human embryo reference^34^ summarizing embryonic stage of the datapoints (generated using https://petropoulos-lanner-labs.clintec.ki.se/shinys/app/ShinyEmbryoRef), E-feature plots of expression of VENTX, GBX2, CR1L and IRX4 genes in the integrated human embryo reference^34^ (generated using https://petropoulos-lanner-labs.clintec.ki.se/shinys/app/ShinyEmbryoRef), F-heatmap of z-score of TPM expression values of the 150 genes upregulated in formative pluripotent stem cell population as defined in^78^, in CHR and parental iPSC lines, G - heatmap of z-score of TPM expression values of the 150 genes upregulated in primed pluripotent stem cell population as defined in^78^, in CHR and parental iPSC lines, H-heatmap of z-score of TPM expression values of the 150 genes upregulated in formative pluripotent stem cell population as defined in^78^, in competent and non-competent iPSC lines (expression data from hipsci.org, Table S2), I - heatmap of z-score of TPM expression values of the 150 genes upregulated in primed pluripotent stem cell population as defined in^78^, in competent and non-competent iPSC lines (expression data from hipsci.org, Table S2)

## Materials and methods

### Cell lines

Three human ESC lines, and thirteen human iPSC lines were used in this study. The ESC lines H1 (WA01, male) and H9 (WA09, female) were purchased from WiCell and the ESC line HUES8-iCas9 (RRID:CVCL_VR10, male) was a gift from Danwei Huangfu.^1^ The iPSC lines: aion2 (HPSI0115i-aion_2, hPSCreg WTSIi255-B, male) bimq4 (HPSI0115i-bimq_4, hPSCreg WTSIi727-A), burb1 (HPSI0714i-burb_1, hPSCreg WTSIi257-A, male), cuhk2 (HPSI0314i-cuhk_2, hPSCreg WTSIi077-B, female), deyz2 (HPSI0215i-deyz_2, hPSCreg WTSIi293-B, female), eesb1(HPSI1014i-eesb_1, hPSCreg WTSIi317-A, female), fawm2 (HPSI0215i-fawm_2, hPSCreg WTSIi178-A, femlae), feec2 (HPSI0214i-feec_2, hPSCreg WTSIi135-A, male), fiaj1 (HPSI0514i-fiaj_1, hPSCreg WTSIi301-A male), giju3 (HPSI0214i-giju_3, hPSCreg WTSIi296-B, female), hehd1 (HPSI1213i-hehd_1, hPSCreg WTSIi003-B, female), kolf2 (HPSI0114i-kolf_2, hPSCreg WTSIi018-B, male) and sojd3 (HPSI0314i-sojd_3, hPSCreg WTSIi073-A, female) were obtained from hispci.org. Approval for the use of human ESCs in this project was granted by the U.K. Stem Cell Bank Steering Committee. Both iPSCs and ESCs were approved by an ERC ethics committee and are registered on the Human Pluripotent StemCell Registry (hpscreg.eu).

### Cell culture

All cell lines were cultured under atmospheric oxygen, 5% CO_2_ and at 37°C in Essential 8 (E8) medium (Thermo Fisher A1517001) on TC-treated 6 well plates (Corning 3506) coated with rh-VTN (Thermo Fischer A14700) at 10 μg/ well in PBS without calcium or magnesium. Cultures were split as clumps twice a week using 0.5mM EDTA. Where specified, culture plates were coated with rhL521 (Gibco A29249) at 5μg per well in DPBS with calcium and magnesium (Gibco 14040133). For some experiments cells were cultured in Stemflex (ThermoFisher, A3349401) or in Stemflex supplemented with 3μM IWP2 (Cayman Chemicals 13951).

### Cerebral organoid generation

Cerebral organoids were generated as described previously with STEMdiff Cerebral Organoid Kit (StemCell Technologies, 08570).^2^ In brief, cultures below 80% confluent were washed in PBS and dissociated with Accutase (Sigma A6964-100 ML), then 9,000 cells per well were seeded in ultra-low adhesion 96 well plate (Corning 7007) in EB medium with 50μM Y27632 (Santa Cruz sc-281642A). After 3 days the formed embryoid bodies were fed with fresh EB medium without Y27632 and then changed to NI on day 5. On day 7 EBs were embedded in Matrigel (Corning 356235) droplets and transferred to Expansion Medium, and then to Maturation Medium on day 10. On day 13 Matrigel was manually dissected and/or dissolved for 30 minutes in Cell Recovery Solution (Corning 354253), and tissues were transferred to fresh Maturation Medium and cultured with agitation.

### Histological and immunohistochemical analysis

Tissues were fixed in 4% PFA either at room temperature for 1h or overnight at 4 °C, then washed twice in PBS for 10 min. Samples for cryo-sectioning were sunk overnight in 30% sucrose in 0.2M PB (21.8 g/l Na2HPO4, 6.4 g/l NaH2PO4 in dH2O), embedded in gelatine (7.5% gelatine, 10% sucrose in 0.2 M PB) and frozen in 2-methylbutane (Sigma-Aldrich, M32631) cooled down to below -30 °C. Gelatine blocks were sectioned at 20 μm and stained as previously described. ^3^ Wholemount samples were stained for 24-48h with primary antibodies followed by 24-48 h with secondary antibodies in a buffer of 4% normal donkey serum and 0.25% Triton-X-100 in PBS.

### Antibodies and stains

SOX2 (Abcam, ab97959, 1:200 for IF), SOX2 (R&D, AF2018, 1:200 IF), SOX2 (Abcam, ab79351, 1:200 for IF), 1:200 TBR2 (R&D Systems, AF6166 1:200 for IF), TBR2 (Abcam, ab216870, 1:200 for IF), SOX10 (R&D Systems, AF2864, 1:500 for IF), SOX17 (R&D Systems, AF1924 1:200 for IF), CDX2 (Cell Signalling Technology, 12306, 1:200 for IF), Villin1 (Proteintech, 16488-1-AP-150UL, 1:100 for IF), ECAD (BD Transduction, 610181, 1:400 for IF), PAX8 (Abcam, ab189249, 1:200 for IF), TCF-2/HNF-1 beta ((R&D Systems, AF3330 1:200 for IF), OCT4 (Abcam, ab181557, 1:200 for IF), Nanog (Abcam, ab21624, 1:100 for IF).

Secondary antibodies used were: Donkey a-Ms 488 (Life Technologies, A21202), Donkey a-Rb 488 (Life Technologies, A21206), Donkey a-Sh 488 (Life Technologies, A11015), Donkey a-Ms 568 (Life Technologies, A10037), Donkey a-Rb 568 (Life Technologies, A10042), Donkey a-Ms 647 (Life Technologies, A31571), Donkey a-Rb 647 (Life Technologies, A31573), Donkey a-Sh 647 (Life Technologies, A21448). All secondary antibodies were used at 1:500 dilution. Nuclei were counterstained with 0.1 µg/ml DAPI (Merck, 268298).

### Imaging and image analysis

Stained samples were imaged on Zeiss LSM 710 or Zeiss LSM 780 systems. Each channel was captured as a separate track. Images were acquired at x100 or x200 magnification. Raw images were processed using FIJI. Brightness and/or contrast were adjusted where needed for visual clarity.

### Chromatin restoration (CHR)

All cultures were carried out under atmospheric oxygen, 5% CO_2_ and 37°C. Human pluripotent stem cells were split using EDTA from their usual culturing conditions and seeded on rhL521-coated plates in Stemflex. When cultures reached 50-60% confluence, culture medium was switched to Medium 1 and cultures were fed daily for 5 days. Medium 1 consisted of KnockOut DMEM (Thermo Fisher 10829018) supplemented with 1% N2 (Thermo Fisher 17502048), 2% B27 (Thermo Fisher A3582801), 1% NEAA (Thermo Fisher 11140050), 1% Glutamax (Thermo Fisher 35050061), 1% Penicillin-Streptomycin (Thermo Fisher 15140122), 50 μg ml^−1^ L-Ascorbic acid 2-phosphate sesquimagnesium salt hydrate (Merck A8960), 20 ng ml^−1^ heregulin beta-1 (Thermo Fisher 100-03-50UG), 1 μM CHRI99021 (Tocris 4423), 2.5 μM IWP2, 10 μM Y27632 dihydrochloride, 0.5 μM SB590885 (AdipoGen Life Sciences SYN-1077), 1 μM PD0325901 (Cell Guidance Systems SM26-2), 500 μM Valproic acid (Merck PHR1061), 5 μM Tranylcypromine (Cayman Chemicals 10010494), 50 nM DZNep (Apexbio A8182), 5 μM EPZ004777 (Cayman 16173) and 1 μM BIX01294 (Santa Cruz Biotechnology sc-293525). From day 6 cultures were fed daily with Medium 2 consisiting of KnockOut DMEM supplemented with 1% N2, 2% B27, 1% NEAA, 1% Glutamax, 1% Penicillin-Streptomycin, 50 μg ml^−1^ L-Ascorbic acid 2-phosphate sesquimagnesium salt hydrate, 20 ng ml^−1^ heregulin beta-1, 1 μM CHRI99021, 2.5 μM IWP2, 10 μM Y-27632, 0.5 μM SB590885 and 1 μM PD0325901. Finally, 5-14 days later, medium was switched to Medium 3 consisting of KnockOut DMEM supplemented with 1% N2, 2% B27, 1% NEAA, 1% Glutamax, 1% Penicillin-Streptomycin, 50 μg ml^−1^ L-Ascorbic acid 2-phosphate sesquimagnesium salt hydrate, 20 ng ml^−1^ heregulin beta-1, 1 μM CHRI99021, 2.5 μM IWP2, 10 μM Y-27632, 0.5 μM SB590885, 0.5 μM PD0325901, 100ng ml^−1^ bFGF (Peprotech 100-18B) and 100ng ml^−1^ mg AlbuMAX™ II Lipid-Rich BSA (Thermo Fisher 11021029). Medium was changed daily for up to 10 days or until colonies with morphology reminiscent of human primed pluripotent cell emerged. These were then picked with a p200 tip and transfereed onto fresh rhL521 coated plates with Stemflex with 2-3 μM IWP2 and 10 μM Y-27632 dihydrochloride. After 24h Y-27632 was dropped and medium was refreshed daily. Alternatively, cultures straight from Medium 3 were switched to Stemflex with 2-3 μM IWP2 and emerging colonies were picked in the same way.

In some instances cultures were passaged when grown in Medium2 or 3 but generally passaging was avoided. To passage the cells, the cultures were washed with PBS without calcium or magenesium, incubated with Accutase (Thermo Fisher A1110501) for 4 min and then then re-suspended in double the volume of medium. Cell suspension was then spun down at 300rcf for 2 min and cell pellet re-suspended in fresh medium. Cultured were passaged at the ratio of 1:2 or 1:3 by surface area.

Established cultures in Stemflex IWP2 were then expanded on rhL521 in Stemflex 2-3 IWP2 and used for experiments or cryopreserved.

### Intestinal organoid generation

Intestinal organoids were generated according to published protocol with the following modifications: cells were aggregated in Aggrewell instead of spontaneous budding and epiregulin was used instead of EGF. ^96-98^ Briefly, human pluripotent stem cells were seeded at 150,000 cell per well of a 6 well plate and cultured for 72hrs in Semflex on Matrigel (Corning 356230, 0.083mg/ml in DMEM/F12 (Thermo Fisher 11320033)) coated plates whereby they achieved 80-90% confluence. The cells were then switched to basal induction medium (RPMI 1640 (Gibco 61870-010), 2% B-27 and Pen/Strep (Gibco 15140148)) with the addition of 100ng/ml Activin A (Invitrogen 17851733) (Definitive Endoderm induction medium) for definitive endoderm differentiation (Day1) and fed daily for two more days. On Day 4, the medium was replaced with hindgut induction medium (basal induction medium with 2μM CHIR and 500ng/ml FGF4(R&D Systems 7460-F4)) and refreshed daily. On day 8 the cultures were dissociated with Accutase and seeded on Aggrewell-400 24-well plates (Stemcell Technologies, 34411) at 2x10^5^ cells per well (500 cell per aggregate). On day 9, the aggregates were washed out of Aggrewells and resuspended in a dome of Matrigel, which was then placed in 6 well suspension plate (Sarstedt 83.3920.500) at the ratio of 1 well in Aggrewell 400 to 1 well in 6 well suspension plate. Matrigel was left to polymeryse for 20 minutes at 37°C and then 2ml of intestinal differentiation medium (intDM) was gently overlayed on top. The intDM comprised Advanced DMEM/F12 (Gibco 12634-010) with 2% B-27, 1% Pen/Strep, 2% Glutamax and 15 μM HEPES (Gibco 15630-049) supplemented with 10ng/ml Epiregulin (Peprotech 100-04), 500ng/ml Rspondin (R&D Systems 4645-RS-025) and 100ng/ml noggin (Peprotech 10-10C). From day 12 Rspondin and noggin were dropped, and medium was changed twice a week. Intestinal organoids were fixed on day 30 in 4% PFA for 1 hour, washed in PBS and then processed for immunofluorescence.

### Kidney organoid generation

Kidney organoids were generated according to published protocol with minor modifications.^72^ Briefly, 96,000 human pluripotent stem cells were seeded on Geltrex (Thermo Fisher A14133-01)-coated plates in Stemflex with 10 µM ROCK inhibitor (Y27632) and after 24hours the medium was changed to TeSR-E6 (Stemcell Technologies, 05946) with 6μM CHIR99021 (Day0). On Day 4, medium was changed to TeSR-E6 with 1μM CHIR and 200ng/ml FGF9 (PeproTech, AF-100-23). On Day 7, the induced cells were dissociated with TrypLE (Thermo Fisher 12604013) and seeded into Aggrewell-400 24-well plates at 1.2x10^6^ cells per well (24-well) in TeSR-E6 with 1μM CHIR and 200ng/ml FGF9. Plates were spun down at 200g for 5 mins, and organoids left to form at 37°C with 5% CO_2_. On Day 10, the organoids from each well were transferred into 6-well ultra-low-adhesion plates (Greiner Bio-One, 657185) in fresh TeSR-E6 with 1μM CHIR and 200ng/ml FGF9, and incubated with agitation at 37°C and 5% CO_2_. On Day 12, medium was replaced with TeSR-E6. Organoids were analysed on day 13 or 15 by phase contrast microscopy or fixed in 4% PFA for 20 minutes and then processed for immunofluorescent staining.

### ATAC-seq

ATAC-seq was performed using the Active Motif ATAC-seq kit (Active Motif 53150).^99^ Briefly, Adherent cells were treated with Accutase to produce a single cell suspension. 100,000 cells per sample were pelleted, washed once in ice-cold PBS, resuspended in ATAC Lysis Buffer and then combined with Tagmentation Master Mix. The reaction was then incubated for 30min at 37°C with 800rpm agitation. Immediately after, the tagmented DNA was column-purified, eluted in a 35μL volume and PCR-amplified for 10 cycles with indexing primers to generate sequencing libraries. Library quality control was carried out with the Bioanalyzer High-Sensitivity DNA analysis kit (Agilent 5067-4626). Libraries were sequenced as paired end 50bp reads, on the Illumina NovaSeq platform at the Cancer Research UK Cambridge Institute Genomics Core Facility. Two biological replicates were collected from independent experiments, except from HUES8 and fiaj1 lines in the competent vs non-competent comparison, where only one biological repeat was available.

### ATAC-seq data analysis

Quality control, trimming and mapping of the FASTQ files was performed using the *atacseq* pipeline (version 2.1.2), distributed by *nf-core [*doi: 10.5281/zenodo.2634132]. The pipeline was configured so that each constituent component was under version control and executed within a Singularity container [https://sylabs.io]. The following command was used to execute the pipeline:

nextflow run nf-core/atacseq -r 2.1.2 --input batch2_samplesheet.csv --genome homo_sapiens.GRCh38.release_102 -config /public/singularity/containers/nextflow/lmb-nextflow/lmb.config --read_length 50 --trim_nextseq 20 --mito_name MT --blacklist blacklist/ENCFF356LFX.bed.chr_deleted.txt --aligner bowtie2 --deseq2_vst --outdir results - bg

These parameters generally adhere to the defaults set by the *nf-core atacseq* pipeline, except that Bowtie2 (version 2.4.4)^100^ was used as the aligner for the pipeline. Reads were mapped to the human reference genome (GRCh38 / release 102), but reads mapping to the mitochondrial chromosome or regions blacklisted by the ENCODE project were removed [https://www.encodeproject.org/files/ENCFF356LFX/]. The specified read trimming parameters (--trim_nextseq 20) were sequencing platform-specific, to account for the Illumina colour chemistry in which high quality calls for guanine nucleotides may actually reflect no signal on a flowcell. The pipeline was also passed a sample list detailing the forward and reverse FASTQ file pairings, along with the sample identities and how technical and biological replicates should be grouped together.

The *nf-core atacseq* pipeline used the software *MACS2* (version 2.2.7.1)^101^ to identify peaks i.e genomic regions with a localised high abundance of mapped reads. MACS2 was run using the default settings for the *atacseq* pipeline. Peaks present in at least 2 of any of the biological replicates were considered as “consensus peaks” and retained. Since the start and end positions of peaks rarely match to the exact base position between samples, overlapping peak regions from different biological replicates were merged together.

The number of reads mapping to each peak was quantitated using the genome browser and analysis software SeqMonk (version 1.48.1) [https://www.bioinformatics.babraham.ac.uk/projects/seqmonk]. To achieve this, BAM files of the aligned reads generated by the *nf-core atacseq* pipeline were imported into SeqMonk, along with a text-format list detailing ATAC-seq peak genomic locations. By using the “Feature Probe Generator” and “Read Count Quantitation” SeqMonk functions, an “Annotated Probe Report” was created listing the raw read counts for each peak. A separate file was also created using this method, but instead it reported the log2-transformed normalised data (counts per million reads) for each peak. A minimum threshold of -2 (log2-transformed counts per million reads) in any of the datasets was applied to remove consensus peaks with low count values.

Principal component analysis of the quantitated peak data was performed using the sklearn.decomposition.PCA class from scikit-learn. ^102^ The input data was the log2-transformed counts per million quantitated peak data generated using SeqMonk.

Classifying the peaks based on their genomic location (e.g. whether they were positioned in genic or intergenic regions) was achieved using the software package Homer (v5.0.1). ^103^ Specifically, the Perl script *annotatePeaks.pl* was run taking ATAC-seq peak co-ordinates BED files and a human genome (GRCh38) GTF annotation file as input.

Raw counts of reads mapping to peak regions for each biological replicate were generated by SeqMonk in the form of an “Annotated Probe Report”. Differential peaks were then identified using the Bioconductor package DESeq2 (version 1.42.1)^104^ running in R (version 4.3.3) *[R Core Team, 2024, R: A Language and Environment for Statistical Computing,* https://www.R-project.org].

DESeq2 was used to identify peaks in which their degree of openness corresponded with cell competence. The count data from the raw biological replicates of competent (H1, H9, hehd1 and kolf2) and not competent (HUES8, fiaj1, sojd3) cell lines was used as the input.

Cell line competence and the sex of the cell line donor were used as the 2 terms for the DESeq2 design formula, with the cell line competence specified as the variable of interest. Similarly, differential peaks before and after CHR were identified in the cell lines fiaj1 and sojd3. Cell line identity and whether cells had undergone treatment were used as the 2 terms for the DESeq2 design formula, with treatment specified as the variable of interest. Differential peaks were defined as having a minimum absolute log2-fold change of 1 and a maximum adjusted p-value of 0.05.

The metagene profile plots were generated using SeqMonk (version 1.48.1). The software generated 500bp tiled windows across the genome. The number of reads mapping to each bin was subsequently quantitated and normalised by library size (i.e. dividing by the total number of reads in each library and then multiplying by 1 million). The SeqMonk “Quantitation Trend Plot” function was used to calculate the normalised read count across each gene, in addition to flanking regions 5kb up- and downstream. The average values for all genes at each given position were then displayed on the metagene plot.

The read distribution heatmaps were generated using SeqMonk (version 1.48.1). The “Aligned Probes Plot” function of the software was used to calculate the intensity of reads across peaks and their associated up- and downstream flanking 2kb regions. The datasets were normalised for total library size and each dataset was clustered independently. The tool “Analysis of Motif Enrichment” (version 5.5.5)^105^ was used to identify known motifs that were relatively enriched in the differentially changing peaks.

The AME package searched the JASPAR CORE (non-redundant, vertebrate, 2022) motif database^106^ to identify previously validated motif sequences.

The total consensus peaks were used as a control background list of sequences for this analysis i.e. motifs needed to be enriched in the peaks of interest as compared to the consensus ATAC-seq peaks.

The enrichment model of the AME software required that the peak length distributions of the target list and the control list should approximately correspond to each other. Since this was not the case, the control list was down-sampled to match the target list in this regard. This was achieved by fitting both the target and control peak lengths to separate Beta distributions. Using these Beta distributions, peaks could be randomly selected from control peaks in a fashion weighted by length. Consequently, the final down-sampled, control list, peak length distribution then matched the peak length distribution of the target list.

The classification of the transcription factors for which binding motifs were enriched was annotated manually using http://tfclass.bioinf.med.uni-goettingen.de.

### Bulk RNAseq sample preparation

700,000 human induced pluripotent stem cells (hiPSCs) or human embryonic stem cells (hESCs) were used for RNA extraction for each biological replicate. Cells were washed once with 1ml Dulbecco’s Phosphate Buffered Saline (DPBS) and harvested using 500µl Accutase (StemCell Technologies, 07922) for 3 minutes at 37°C. Dissociated cells were then collected with 5ml of corresponding PSC medium and centrifuged for 3 min at 400 RCF at RT. Supernatant was aspirated and the cells were resuspended in 1ml cold DPBS.

RNA extraction from hiPSCs/hESCs was conducted using the Monarch Total RNA Miniprep Kit (NEB T2010S) according to the manufacturer’s protocol, with DNase I treatment used to enzymatically remove residual gDNA. RNA concentration was quantified using Nanodrop and the RNA Integrity Number (RIN) was assessed using the Agilent RNA 6000 Pico Kit (Agilent, 5067-1513) on Agilent 2100 Bioanalyzer (Agilent). A RIN value of >7 was deemed to be of good quality and used for downstream RNA-seq library processing. The isolated RNA was frozen and stored at -80°C until RNA-seq library preparation.

RNA-seq library preparation was performed on the isolated RNA using the NEBNext Poly(A) mRNA Magnetic Isolation Module, which isolated mRNA from total RNA (100ng input) by poly(A)-enrichment. RNA-seq libraries were subsequently prepared from the isolated mRNA using the NEBNext Ultra II Directional RNA Library Prep Kit from Illumina, the NEBNext Poly(A) mRNA Magnetic Isolation Module (NEB, E7490L), and the NEBNext Multiplex Oligos from Illumina (Index Primers Set 1, 2, 3 and 4, NEB, E7335S, E7500S, E7710S, E7730S), following the manufacturer’s protocols. The final sequencing-ready libraries were amplified using the protocol’s PCR programme with 12 cycles. Size distribution and yield quantification of the RNA-seq libraries were assessed using an Agilent 2100 Bioanalyzer high sensitivity DNA chip (Agilent 5067–4626). Libraries were then multiplexed and sequenced (paired-end 150 bp) on a NovaSeq X Plus Series PE150 (Illumina) to a depth of a minimum 30 million reads per library.

### Bulk RNAseq analysis

Quality control, trimming and mapping of the FASTQ files was performed using the *rnaseq* pipeline (version 3.12.0), distributed by *nf-core [doi:* https://zenodo.org/records/7998767]. The pipeline was configured so that each constituent component was under version control and executed within a Singularity container [https://sylabs.io]. The following command was used to execute the pipeline:

nextflow run nf-core/rnaseq -r 3.12.0 --input samplesheet.csv --genome homo_sapiens.GRCh38.release_102 -config config.txt --extra_trimgalore_args ’--nextseq 20’ --outdir results --deseq2_vst -bg

These parameters generally adhere to the defaults set by the *nf-core rnaseq* pipeline. That pipeline release used STAR (version 2.7.9a)^107^ as the aligner, mapping reads to the human reference genome (GRCh38 / release 102).

The specified read trimming parameter (--nextseq 20) were sequencing platform-specific, to account for the Illumina colour chemistry in which high quality calls for guanine nucleotides may actually reflect no signal on a flow cell. The pipeline was also passed a sample list detailing the forward and reverse FASTQ file pairings, along with the sample identities and how any technical and biological replicates should be grouped together.

The *nf-core rnaseq* pipeline produces quantitated expression values for every gene for each sample, reporting raw counts and TPM values. These expression matrices were used in subsequent analyses.

Raw read counts per gene were generated by the *nf-core rnaseq* pipeline for each biological replicate. These data were subsequently normalised using the Variance Stabilising Transformation algorithm of DESeq2 (version 1.42.1)^104^ running in R (version 4.3.3). The normalised data were filtered, using the VST threshold of 7 and used for principal component analysis. Principal component analysis of the normalised data was performed using the *sklearn.decomposition.PCA* class from scikit-learn. ^102^

Raw read counts per gene were generated by the *nf-core rnaseq* pipeline for each biological replicate. Differentially expressed genes were then identified using the Bioconductor package DESeq2 (version 1.42.1)^104^, running in R (version 4.3.3) *[R Core Team, 2024, R: A Language and Environment for Statistical Computing,* https://www.R-project.org]. Differential genes were defined as having a minimum absolute fold change of 1.5 and a maximum adjusted p-value of 0.05, as calculated by DESeq2.

### Single cell RNA-seq analysis

*Pooled iPSC cerebral organoids*

Single cell RNA-seq data from two batches of organoids generated from a pool of 18 iPS cells were previously published^16^ and deposited on EBI-ENA, study accession PRJEB38269. Run accessions for batch 1 were ERR470047, ERR4700472, ERR4700473, ERR4700474 and for batch 2 were ERR4700475, ERR4700476, ERR4700477. The data were mapped as described in the single cell RNA-seq mapping section below, except that the reads were mapped to GRCh37. This previous genome annotation was selected to match the Variant Call Format (VCF) files obtained from the Human Induced Pluripotent Stem Cells Initiative (HipSci) [https://www.hipsci.org]. Using these variant calls for each cell line, in conjunction with the mapped sequence data, enabled the donor of origin to be ascertained for individual cells. More specifically, this donor-level classification of cells was performed using the software Demuxlet^108^ The code was obtained from the Demuxlet GitHub repository [https://github.com/statgen/demuxlet] in March 2024 and was run with default settings Only Demuxlet classified single cell barcodes were carried forward, and doublets or ambiguous called cell barcodes were excluded from subsequent analysis.

For the single cell analysis pipeline the reads were mapped onto Human Genome Reference 38 using Cell Ranger^109^. The expression matrix was then analysed in Seurat v5.0.3 and low-quality cells with percent.MT >20 were filtered out. Seurat v5 integration was performed for batch correction using CCAIntegration, followed by FindNeighbors, FindClusters resolution.2, and RunUMAP dimensions 1:15. Clusters were annotated based on the top 30 differential genes and a panel of well-described markers: SOX2, HOPX (radial glia); EOMES, NEUROD4 (INPs); NEUROD2, DCX (neurons); SATB2 (UL neurons); BCL11B (DL neurons; TTR, RSPO2 (hem/ChP); FN1, TGFB1, PDGFRA, LUM (fibroblasts); PAX7, MYF5 (satellite cells); MYOD1, TTN (muscle); CENPF, MKI67 (cycling cells).

### Stem cell embryo model

The single cell RNA-seq (10x Genomics Chromium v3.1 Dual Index system) dataset SRX19950187 was obtained from the National Center for Biotechnology Information [https://www.ncbi.nlm.nih.gov/sra?term=SRX19950187].

Quality control and mapping of the FASTQ files was performed using the single cell RNA-seq pipeline *scrnaseq* (version 2.5.1), distributed by *nf-core [doi:* https://zenodo.org/records/10554425]. The pipeline was configured so that each constituent component was under version control and executed within a Singularity container [https://sylabs.io]. The following command was used to execute the pipeline:

nextflow run nf-core/scrnaseq -r 2.5.1 --input samplesheet.csv --genome homo_sapiens.GRCh38.release_102 --aligner cellranger -config config.txt --outdir results -bg These parameters generally adhere to the defaults set by the *nf-core scrnaseq* pipeline, except that Cell Ranger (version 7.1.0)^109^ was used as the aligner. Reads were mapped to the human reference genome (GRCh38 / release 102). Cell Ranger generated a filtered HDF5 feature barcode matrix object, which was imported into an AnnData object using Scanpy^110^ (version 1.9.8).

The imported dataset underwent quality control. Specifically, cells in which more than 15% of the reads mapped to the mitochondrial chromosome were removed, as were cells in which fewer than 1,000 genes exhibited detectable expression. Conversely, cells with more than 10,000 expressed genes were also removed from subsequent analysis. Genes whose transcripts were observed in fewer than three cells were also deleted from the dataset. The software package Scrublet (version 0.2.3) was used to remove putative technical doublets (formed by the random co-encapsulation of two cells).

The read count for each cell was normalised by total counts over all genes, so that every cell had the same total count after normalisation. The dataset was then log(x+1) transformed. The top 2,000 highly variable genes were identified and used to visualise the relatedness of different cells in the form of a UMAP. Different cell types were subsequently identified on the UMAP by using a Leiden graph-clustering method, setting the resolution to 1.2. Cellular identities were then determined by ascertaining the expression profile of a panel of pre-selected marker genes for each of these clusters.

### Integrating single cell RNA-seq data with bulk RNA-seq data

Raw read counts for each gene in the pre-determined Leiden clusters were aggregated at the cell level. The resulting expression data matrix was then merged with the expression data for the bulk RNA-seq data. Batch correction was performed using ComBat-seq (SVA version 3.40.0)^111^ in R (version 4.1.0). When performing the batch correction with ComBat-seq, the single cell-derived “pseudo-bulk” data was considered as a distinct batch, as were other independent sequencing runs.

These data were subsequently normalised using the Variance Stabilising Transformation algorithm of DESeq2 (version 1.42.1)^105^ running in R (version 4.3.3). The normalised data were filtered, using the VST threshold of 7 and used for principal component analysis. Principal component analysis of the normalised data was performed using the *sklearn.decomposition.PCA* class from scikit-learn. ^102^

### CUT&Tag

CUT&Tag was carried out as described previously^75^with minor modifications as per the EpiCypher CUTANA™ CUT&Tag Protocol v 1.7. 120,000 cells per CUT&Tag experiment were harvested for nuclei extraction. The cell suspension was centrifuged for 3 minutes at 300 RCF at RT. The cell pellet was resuspended in 100 µl/sample of cold nuclei extraction buffer (20 mM HEPES–KOH, pH 7.9, 10 mM KCl, 0.1% Triton X-100, 20% Glycerol, 0.5 mM Spermidine (Sigma-Aldrich, 05292), 1x Roche cOmplete^TM^, Mini, EDTA-free Protease Inhibitor) and incubated on ice for 10 minutes, followed by centrifugation for 3 minutes at 600 RCF and 4°C. The nuclei pellet was resuspended in nuclei extraction buffer to achieve a final concentration of 1.2 million nuclei/ml and frozen at -80°C until library preparation. Upon thawing of the nuclei suspension, 100µl of the suspension (120,000 nuclei) was aliquoted to be used per CUT&Tag reaction.

Concavalin A (ConA) beads (EpiCypher, 21-1401) were transferred to a 1.5ml Eppendorf tube on a magnetic stand and washed twice with cold bead activation buffer (20 mM HEPES pH 7.9, 10 mM KCl, 1 mM CaCl_2_, 1 mM MnCl_2_). Beads and buffer were added at 11µl and 100µl per sample, respectively, and after washing, the beads were resuspended in a final volume of 11μl of buffer per sample. 10µl of ConA bead slurry was aliquoted into each sample tube of an 8-strip tube containing 100µl nuclei per tube for batch processing. The ConA bead and nuclei mixture was vortexed gently to mix and incubated at RT for 10 minutes. The tubes were placed on a magnetic stand and the supernatant removed. Each nuclei sample was resuspended in 50µl of cold Antibody150 buffer (20 mM HEPES, pH 7.5, 150 mM NaCl, 0.5 mM Spermidine, 1x Roche cOmplete^TM^, Mini, EDTA-free Protease Inhibitor, 0.01% digitonin (Sigma-Aldrich, D141-100MG), 2mM EDTA) containing primary antibody (anti-H3K27me3 (Active Motif; 39157; dilution 1:50), anti-H3K4me3 (Active Motif; 39160; dilution 1:50), anti-IgG negative control antibody (EpiCypher; 13-0042; dilution 1:50). Samples were then gently mixed and incubated overnight at 4°C on an orbital shaker.

The next day, the tubes were placed on a magnet and the supernatant discarded. Each sample was resuspended in 50µl Digitonin150 buffer (20 mM HEPES, pH 7.5, 150 mM NaCl, 0.5 mM Spermidine, 1x Roche cOmplete^TM^, Mini, EDTA-free Protease Inhibitor, 0.01% digitonin) containing 0.5μg of secondary antibody (anti-rabbit Epicypher; 13-0047) for 1 hr at RT on a nutator. After the incubation, the samples were placed on a magnet and washed twice with Digitonin150 buffer. The samples were then resuspended in 50µl Digitonin300 buffer (20 mM HEPES, pH 7.5, 300 mM NaCl, 0.5 mM Spermidine, 1x Roche cOmplete^TM^, Mini, EDTA-free Protease Inhibitor, 0.01% digitonin), and 1.25μl of CUTANA pAG-TN5 (EpiCypher, 15-1017) was added. Samples were then gently mixed and incubated for 1hr at RT on a nutator. After the incubation, the samples were placed on a magnet and washed twice with Digitonin300 buffer (20mM HEPES, pH 7.5, 300mM NaCl, 0.5mM Spermidine, 1x Roche cOmplete^TM^, Mini, EDTA-free Protease Inhibitor, 0.01% digitonin). The samples were then resuspended in 50 µl of Tagmentation buffer (Digitonin 300 Buffer, 10 mM MgCl_2_) and incubated for 1hr at 37°C in a thermocycler. Then the sample tubes were placed on a magnet and the supernatant discarded, the samples were then resuspended in 50µl TAPS buffer (10 mM TAPS, pH 8.5, 0.2 mM EDTA). The tubes were returned to the magnet and the supernatant removed, followed by addition of 5µl SDS release buffer (0.5M HEPES-NaOH pH 8.5, 10% SDS). The reaction mixture was vortexed on maximum speed for 10s and briefly centrifuged. The samples were then incubated on the thermocycler for 1hr at 58℃. The SDS was neutralised by the addition of 15µl SDS quench buffer (0.67% Triton-X 100), vortexed at maximum speed for 10s, and centrifuged briefly.

The libraries were amplified using KAPA HiFi HotStart ReadyMix (Roche Diagnostics Ltd, KK2602). 25ul KAPA HiFi HotStart ReadyMix per CUT&Tag reaction (mastermix only) was incubated at 95℃ for 3 min and cooled to 22℃ in a thermocycler. 1μl of each of barcoded i5 and i7 primers (5μM stock) and 25µl of the previously heated mastermix was added to the DNA. PCR was conducted using the following programme: extension step at 72°C for 5 min, denaturation at 98°C for 45 s, 13-15 cycles (denaturation at 98°C for 10s and short extension at 60°C for 10s), and final extension at 72°C for 1 min. Amplified libraries were cleaned twice using 1X volume of AMPure XP Beads (Beckman Coulter; A63881) and resuspended in 15μl of 0.1X TE buffer.

Size distribution and yield quantification of the CUT&Tag libraries were tested using an Agilent 2100 Bioanalyzer high sensitivity DNA chip (Agilent 5067–4626). Libraries were then multiplexed and sequenced (paired-end 150 bp) by Novogene on a NovaSeq X Plus Series PE150 (Illumina) to a depth of a minimum 10 million reads per replicate.

### Cut&Tag data analysis

Quality control, trimming and mapping of the FASTQ files was performed using the *cutandrun* pipeline (version 3.2.1), distributed by *nf-core [*doi: 10.5281/zenodo.2634132]. The pipeline was configured so that each constituent component was under version control and executed within a Singularity container [https://sylabs.io]. The following command was used to execute the pipeline:

nextflow run nf-core/cutandrun -r 3.2.1 --input samplesheet.csv --genome homo_sapiens.GRCh38.release_102 -config config.config --blacklist ENCFF356LFX.txt -- normalisation_mode CPM --use_control false --skip_heatmaps --outdir results -bg

These parameters generally adhered to the defaults set by the *nf-core cutandrun* pipeline. Reads were mapped to the human reference genome (GRCh38 / release 102), but reads mapping to regions blacklisted by the ENCODE project [https://www.encodeproject.org/files/ENCFF356LFX/] were removed. The pipeline was also passed a sample list detailing the forward and reverse FASTQ file pairings, along with the sample identities, antibodies used, and how technical and biological replicates should be grouped together.

The pipeline used the software *SEACR* (version 1.3)^112^ to identify peaks i.e genomic regions with a localised high abundance of mapped reads. SEACR was run using the default settings for the *cutandrun* pipeline, but no IgG control was provided.

Next, consensus peaks were identified separately for the H3K4me3 and the H3K27me3 antibodies. Peaks present in at least 2 of any of the biological replicates were considered as “consensus peaks” and retained. Since the start and end positions of peaks rarely match to the exact base position between samples, overlapping peak regions from different biological replicates were merged together.

The number of reads mapping to each peak was quantitated using the genome browser and analysis software SeqMonk (version 1.48.1) [https://www.bioinformatics.babraham.ac.uk/projects/seqmonk]. To achieve this, the aligned reads BAM files generated by the *nf-core cutandrun* pipeline were imported into SeqMonk, along with a text-format list detailing peak genomic locations. By using the “Feature Probe Generator” and “Read Count Quantitation” SeqMonk functions, an “Annotated Probe Report” was created listing the raw read counts for each peak. A separate file was also created using this method, but instead it reported the log2-transformed normalised data (counts per million reads) for each peak.

Raw read counts per peak were generated by SeqMonk for each biological replicate. These data were subsequently normalised using the Variance Stabilising Transformation algorithm of DESeq2 (version 1.42.1)^104^, running in R (version 4.3.3). Principal component analysis of the normalised data was performed using the *sklearn.decomposition.PCA* class from scikit-learn. ^102^

Promoter regions were defined as being 2kb upstream of genes, as reported in the genome annotation GRCh38 / release 102. Promoters overlapping both H3K3me3 and H3K27me3 consensus peaks were then extracted. From this list, promoters that contained both H3K3me3 and H3K27me3 marks, within the same cell line, were classified as bivalent. Metagene profiles were generated as per the ATAC-seq description.

### Enzymatic methyl-seq (EM-seq)

hiPSCs or hESCs genomic DNA (gDNA) was extracted from isolated nuclei using the Monarch Spin gDNA Extraction Kit (NEB, #T3010S). gDNA concentration was quantified using a Nanodrop. 200ng of gDNA was fragmented to an average size of ∼300bp using a Covaris E220 Sonicator with the following parameters: temperature 7°C, peak incident power 140W, duty factor 10%, cycles per burst 200, treatment time 80s. The sheared gDNA was then used to prepare EM-seq libraries using the NEBNext Enzymatic Methyl-seq kit (NEB, #E7120S) following the manufacturer’s standard size library protocol. The final sequencing-ready libraries were amplified using the protocol’s PCR programme with 4 cycles.

Size distribution and yield quantification of the libraries were tested using an Agilent 2100 Bioanalyzer high sensitivity DNA chip (Agilent 5067–4626). Libraries were then multiplexed and sequenced (paired-end 150 bp) on a NovaSeq X Plus Series PE150 (Illumina) to a depth of a minimum 30 million reads per library.

### EM-seq data analysis

The quality of the raw EM-seq FASTQ reads was assessed using FastQC (https://www.bioinformatics.babraham.ac.uk/projects/fastqc/). Reads were subsequently trimmed using Trim Galore v0.6.1.10 (with Cutadapt v4.6) and trimmed reads were reassessed using FastQC. The adaptor trimmed reads were aligned and mapped to *Homo sapiens* GRCh38 with Bismark v0.24.2^113^ with modified trimming parameters optimised for EM-seq. The resulting BAM files were deduplicated and methylation calls were extracted using the Bismark deduplication and methylation extractor tools.

Quantitation of DNA methylation datasets was conducted using SeqMonk. The Bismark-generated methylation call coverage files (.cov files) were imported into SeqMonk. Read positions were generated with CpGs by filtering CpGs with a minimum threshold of 1 read count and grouped into windows containing 100 valid CpGs. The bisulphite methylation feature pipeline in SeqMonk was used to quantitate the CpG windows, with a minimum count of 1 read per CpG position and a minimum of 20 CpGs reads per window. This generated mean percentage methylation values for each window. Differential methylation analysis between incompetent and reset incompetent lines was performed using the EdgeR (for/rev) statistical filter in SeqMonk with a p-value<0.05 cut-off after Benjamimi and Hochberg correction with a minimum methylation difference threshold of 25%.

## Cited literature

1. Rostovskaya, M., Andrews, S., Reik, W. & Rugg-Gunn, P. J. Amniogenesis occurs in two independent waves in primates. Cell Stem Cell 29, 744–759.e6 (2022).

2. Zhai, J., et al. Primate gastrulation and early organogenesis at single-cell resolution. Nature 612, 732–738 (2022).

3. Xiao, Z. et al. 3D reconstruction of a gastrulating human embryo. Cell 187, 2855–2874.e19 (2024).

4. Shahbazi, M. N. & Pasque, V. Early human development and stem cell-based human embryo models. Cell Stem Cell 31, 1398–1418 (2024).

5. Hemmati-Brivanlou, A. & Melton, D. Vertebrate Embryonic Cells Will Become Nerve Cells Unless Told Otherwise. Cell 88, 13–17 (1997).

6. Levine, A. J. & Brivanlou, A. H. Proposal of a model of mammalian neural induction. Dev. Biol. 308, 247–256 (2007).

7. Muñoz-Sanjuán, I. & Brivanlou, A. H. Neural induction, the default model and embryonic stem cells. Nat. Rev. Neurosci. 3, 271–280 (2002).

8. Thomson, J. A. et al. Embryonic Stem Cell Lines Derived from Human Blastocysts. Sci. New Ser. 282, 1145–1147 (1998).

9. Lancaster, M. A. et al. Cerebral organoids model human brain development and microcephaly. Nature 501, 373–379 (2013).

10. Osafune, K. et al. Marked differences in differentiation propensity among human embryonic stem cell lines. Nat. Biotechnol. 26, 313–315 (2008).

11. Hu, B.-Y., et al. Neural differentiation of human induced pluripotent stem cells follows developmental principles but with variable potency. Proc. Natl. Acad. Sci. 107, 4335–4340 (2010).

12. Bock, C. et al. Reference Maps of Human ES and iPS Cell Variation Enable High-Throughput Characterization of Pluripotent Cell Lines. Cell 144, 439–452 (2011).

13. Kajiwara, M. et al. Donor-dependent variations in hepatic differentiation from human-induced pluripotent stem cells. Proc. Natl. Acad. Sci. 109, 12538–12543 (2012).

14. Kilpinen, H. et al. Common genetic variation drives molecular heterogeneity in human iPSCs. Nature 546, 370–375 (2017).

15. Cuomo, A. S. E. et al. Single-cell RNA-sequencing of differentiating iPS cells reveals dynamic genetic effects on gene expression. Nat. Commun. 11, 810 (2020).

16. Jerber, J. et al. Population-scale single-cell RNA-seq profiling across dopaminergic neuron differentiation. Nat. Genet. 53, 304–312 (2021).

17. Puigdevall, P., Jerber, J., Danecek, P., Castellano, S. & Kilpinen, H. Somatic mutations alter the differentiation outcomes of iPSC-derived neurons. Cell Genomics 3, 100280 (2023).

18. Nishizawa, M. et al. Epigenetic Variation between Human Induced Pluripotent Stem Cell Lines Is an Indicator of Differentiation Capacity. Cell Stem Cell 19, 341–354 (2016).

19. Buckberry, S. et al. Transient naive reprogramming corrects hiPS cells functionally and epigenetically. Nature 620, 863–872 (2023).

20. Kinoshita, M. et al. Capture of Mouse and Human Stem Cells with Features of Formative Pluripotency. Cell Stem Cell 28, 453–471.e8 (2021).

21. Osteil, P. et al. Dynamics of Wnt activity on the acquisition of ectoderm potency in epiblast stem cells. Development 146, dev172858 (2019).

22. Tsakiridis, A. et al. Distinct Wnt-driven primitive streak-like populations reflect *in vivo* lineage precursors. Development 141, 1209–1221 (2014).

23. Wu, J. et al. An alternative pluripotent state confers interspecies chimaeric competency. Nature 521, 316–321 (2015).

24. Kurek, D. et al. Endogenous WNT signals mediate BMP-induced and spontaneous differentiation of epiblast stem cells and human embryonic stem cells. Stem Cell Rep. 4, 114–128 (2015).

25. Sáez, M. et al. Statistically derived geometrical landscapes capture principles of decision-making dynamics during cell fate transitions. Cell Syst. 13, 12–28.e3 (2022).

26. Metzis, V. et al. Nervous System Regionalization Entails Axial Allocation before Neural Differentiation. Cell 175, 1105–1118.e17 (2018).

27. Barry, C. et al. Species-specific developmental timing is maintained by pluripotent stem cells ex utero. Dev. Biol. 423, 101–110 (2017).

28. Benito-Kwiecinski, S. et al. An early cell shape transition drives evolutionary expansion of the human forebrain. Cell 184, 2084–2102.e19 (2021).

29. Rayon, T. et al. Species-specific pace of development is associated with differences in protein stability. Science 369, eaba7667 (2020).

30. Veenvliet, J. V., Lenne, P.-F., Turner, D. A., Nachman, I. & Trivedi, V. Sculpting with stem cells: how models of embryo development take shape. Development 148, dev192914 (2021).

31. Sutcliffe, M. A. et al. Optimizing PSC culture for generating neural organoids. Preprint at 10.1101/2024.11.20.624444 (2024).

32. Kojima, Y. et al. The Transcriptional and Functional Properties of Mouse Epiblast Stem Cells Resemble the Anterior Primitive Streak. Cell Stem Cell 14, 107–120 (2014).

33. Oldak, B. et al. Complete human day 14 post-implantation embryo models from naive ES cells. Nature (2023) doi:10.1038/s41586-023-06604-5.

34. Zhao, C. et al. A comprehensive human embryo reference tool using single-cell RNA-sequencing data. Nat. Methods (2024) doi:10.1038/s41592-024-02493-2.

35. Huelsken, J. et al. Requirement for ␤-Catenin in Anterior-Posterior Axis Formation in Mice. J. Cell Biol. 148, (2000).

36. Beddington, R. S. P. & Robertson, E. J. Axis Development and Early Asymmetry in Mammals. Cell 96, 195–209 (1999).

37. Martyn, I., Brivanlou, A. H. & Siggia, E. D. A wave of WNT signaling balanced by secreted inhibitors controls primitive streak formation in micropattern colonies of human embryonic stem cells. Dev. Camb. Engl. 146, dev172791 (2019).

38. Blauwkamp, T. A., Nigam, S., Ardehali, R., Weissman, I. L. & Nusse, R. Endogenous Wnt signalling in human embryonic stem cells generates an equilibrium of distinct lineage-specified progenitors. Nat. Commun. 3, 1070 (2012).

39. Strano, A., Tuck, E., Stubbs, V. E. & Livesey, F. J. Variable Outcomes in Neural Differentiation of Human PSCs Arise from Intrinsic Differences in Developmental Signaling Pathways. Cell Rep. 31, 107732 (2020).

40. Huggins, I. J. et al. The WNT target SP5 negatively regulates WNT transcriptional programs in human pluripotent stem cells. Nat. Commun. 8, 1034 (2017).

41. Burrill, D. R. & Silver, P. A. Making Cellular Memories. Cell 140, 13–18 (2010).

42. Veres, T. et al. Cellular forgetting, desensitisation, stress and ageing in signalling networks. When do cells refuse to learn more? Cell. Mol. Life Sci. 81, 97 (2024).

43. Yoney, A. et al. WNT signaling memory is required for ACTIVIN to function as a morphogen in human gastruloids. eLife 7, e38279 (2018).

44. Yoney, A., Bai, L., Brivanlou, A. H. & Siggia, E. D. Mechanisms underlying WNT-mediated priming of human embryonic stem cells. Development 149, dev200335 (2022).

45. Espinosa-Martínez, M., Alcázar-Fabra, M. & Landeira, D. The molecular basis of cell memory in mammals: The epigenetic cycle. Sci. Adv. 10, eadl3188 (2024).

46. Stewart-Morgan, K. R., Reverón-Gómez, N. & Groth, A. Transcription Restart Establishes Chromatin Accessibility after DNA Replication. Mol. Cell 75, 284–297.e6 (2019).

47. Chen, Z. et al. Increased enhancer–promoter interactions during developmental enhancer activation in mammals. Nat. Genet. 56, 675–685 (2024).

48. Zylicz, J. J. et al. Chromatin dynamics and the role of G9a in gene regulation and enhancer silencing during early mouse development. eLife 4, e09571 (2015).

49. Nord, A. S. et al. Rapid and Pervasive Changes in Genome-wide Enhancer Usage during Mammalian Development. Cell 155, 1521–1531 (2013).

50. Argelaguet, R. et al. Multi-omics profiling of mouse gastrulation at single-cell resolution. Nature 576, 487–491 (2019).

51. Tosic, J. et al. Eomes and Brachyury control pluripotency exit and germ-layer segregation by changing the chromatin state. Nat. Cell Biol. 21, 1518–1531 (2019).

52. Gifford, C. A. et al. Transcriptional and Epigenetic Dynamics during Specification of Human Embryonic Stem Cells. Cell 153, 1149–1163 (2013).

53. Tsaytler, P., Blaess, G., Scholze-Wittler, M., Koch, F. & Herrmann, B. G. Early neural specification of stem cells is mediated by a set of SOX2-dependent neural-associated enhancers. Stem Cell Rep. 19, 618–628 (2024).

54. Blanco, M. A. et al. Chromatin-state barriers enforce an irreversible mammalian cell fate decision. Cell Rep. 37, 109967 (2021).

55. Ciceri, G. et al. An epigenetic barrier sets the timing of human neuronal maturation. Nature 626, 881–890 (2024).

56. Lando, D. et al. Enhancer-promoter interactions are reconfigured through the formation of long-range multiway hubs as mouse ES cells exit pluripotency. Mol. Cell 84, 1406–1421.e8 (2024).

57. Nashun, B., Hill, P. W. & Hajkova, P. Reprogramming of cell fate: epigenetic memory and the erasure of memories past. EMBO J. 34, 1296–1308 (2015).

58. Chen, M. et al. Chromatin architecture reorganization in murine somatic cell nuclear transfer embryos. Nat. Commun. 11, 1813 (2020).

59. Creyghton, M. P. et al. Histone H3K27ac separates active from poised enhancers and predicts developmental state. Proc. Natl. Acad. Sci. 107, 21931–21936 (2010).

60. Gafni, O. et al. Derivation of novel human ground state naive pluripotent stem cells. Nature 504, 282–286 (2013).

61. Buecker, C. et al. Reorganization of Enhancer Patterns in Transition from Naive to Primed Pluripotency. Cell Stem Cell 14, 838–853 (2014).

62. Guo, G. et al. Correction: Epigenetic resetting of human pluripotency (doi:10.1242/dev.146811). *Development* **145**, dev166397 (2018).

63. Guan, J. et al. Chemical reprogramming of human somatic cells to pluripotent stem cells. Nature 605, 325–331 (2022).

64. Xiang, Y. et al. Epigenomic analysis of gastrulation identifies a unique chromatin state for primed pluripotency. Nat. Genet. 52, 95–105 (2020).

65. De Clerck, L. et al. Untargeted histone profiling during naive conversion uncovers conserved modification markers between mouse and human. Sci. Rep. 9, 17240 (2019).

66. Battle, S. L. et al. Enhancer Chromatin and 3D Genome Architecture Changes from Naive to Primed Human Embryonic Stem Cell States. Stem Cell Rep. 12, 1129–1144 (2019).

67. Grosswendt, S. et al. Epigenetic regulator function through mouse gastrulation. Nature 584, 102–108 (2020).

68. Nicetto, D. et al. H3K9me3-heterochromatin loss at protein-coding genes enables developmental lineage specification. Science 363, 294–297 (2019).

69. Zhu, J. et al. Genome-wide chromatin state transitions associated with developmental and environmental cues. Cell 152, 642–654 (2013).

70. Padeken, J., Methot, S. P. & Gasser, S. M. Establishment of H3K9-methylated heterochromatin and its functions in tissue differentiation and maintenance. Nat. Rev. Mol. Cell Biol. 23, 623–640 (2022).

71. Chen, J. et al. H3K9 methylation is a barrier during somatic cell reprogramming into iPSCs. Nat. Genet. 45, 34–42 (2013).

72. Kumar, S. V. et al. Kidney micro-organoids in suspension culture as a scalable source of human pluripotent stem cell-derived kidney cells. Development 146, dev172361 (2019).

73. Spence, J. R. et al. Directed differentiation of human pluripotent stem cells into intestinal tissue in vitro. Nature 470, 105–109 (2011).

74. Bernstein, B. E. et al. A Bivalent Chromatin Structure Marks Key Developmental Genes in Embryonic Stem Cells. Cell 125, 315–326 (2006).

75. Kaya-Okur, H. S. et al. CUT&Tag for efficient epigenomic profiling of small samples and single cells. Nat. Commun. 10, 1930 (2019).

76. Schulz, M. et al. DNA methylation restricts coordinated germline and neural fates in embryonic stem cell differentiation. Nat. Struct. Mol. Biol. 31, 102–114 (2024).

77. Sasaki, K. et al. Robust In Vitro Induction of Human Germ Cell Fate from Pluripotent Stem Cells. Cell Stem Cell 17, 178–194 (2015).

78. Arthur, T. D. et al. Complex regulatory networks influence pluripotent cell state transitions in human iPSCs. Nat. Commun. 15, 1664 (2024).

79. Polo, J. M. et al. A Molecular Roadmap of Reprogramming Somatic Cells into iPS Cells. Cell 151, 1617–1632 (2012).

80. Bertero, A. et al. Activin/Nodal signaling and NANOG orchestrate human embryonic stem cell fate decisions by controlling the H3K4me3 chromatin mark. Genes Dev. 29, 702–717 (2015).

81. Vermaak, D. Maintenance of chromatin states: an open-and-shut case. Curr. Opin. Cell Biol. 15, 266–274 (2003).

82. Gilbert, N. et al. Chromatin Architecture of the Human Genome: Gene-Rich Domains Are Enriched in Open Chromatin Fibers.

83. Tu, Z. et al. Discordance between chromatin accessibility and transcriptional activity during the human primed-to-naïve pluripotency transition process. Cell Regen. 12, 35 (2023).

84. Kiani, K., Sanford, E. M., Goyal, Y. & Raj, A. Changes in chromatin accessibility are not concordant with transcriptional changes for single-factor perturbations. Mol. Syst. Biol. 18, e10979 (2022).

85. Chereji, R. V., Eriksson, P. R., Ocampo, J., Prajapati, H. K. & Clark, D. J. Accessibility of promoter DNA is not the primary determinant of chromatin-mediated gene regulation. Genome Res. 29, 1985–1995 (2019).

86. Carcamo-Orive, I. et al. Analysis of Transcriptional Variability in a Large Human iPSC Library Reveals Genetic and Non-genetic Determinants of Heterogeneity. Cell Stem Cell 20, 518–532.e9 (2017).

87. Holoch, D. et al. A cis-acting mechanism mediates transcriptional memory at Polycomb target genes in mammals. Nat. Genet. 53, 1686–1697 (2021).

88. Sasaki, K. et al. The Germ Cell Fate of Cynomolgus Monkeys Is Specified in the Nascent Amnion. Dev. Cell 39, 169–185 (2016).

89. Chen, D. et al. Human Primordial Germ Cells Are Specified from Lineage-Primed Progenitors. Cell Rep. 29, 4568–4582.e5 (2019).

90. Henrique, D., Abranches, E., Verrier, L. & Storey, K. G. Neuromesodermal progenitors and the making of the spinal cord. Development 142, 2864–2875 (2015).

91. Tien, C.-L. et al. Snail2/Slug cooperates with Polycomb repressive complex 2 (PRC2) to regulate neural crest development. Development dev.111997 (2015) doi:10.1242/dev.111997.

92. Shan, Y. et al. PRC2 specifies ectoderm lineages and maintains pluripotency in primed but not naïve ESCs. Nat. Commun. 8, 672 (2017).

93. Zhao, Y. et al. Coordination of EZH2 and SOX2 specifies human neural fate decision. Cell Regen. 10, 30 (2021).

94. González, F. et al. An iCRISPR Platform for Rapid, Multiplexable, and Inducible Genome Editing in Human Pluripotent Stem Cells. Cell Stem Cell 15, 215–226 (2014).

95. Lancaster, M. A. et al. Guided self-organization and cortical plate formation in human brain organoids. Nat. Biotechnol. 35, 659–666 (2017).

96. McCracken, K. W., Howell, J. C., Wells, J. M. & Spence, J. R. Generating human intestinal tissue from pluripotent stem cells in vitro. Nat. Protoc. 6, 1920–1928 (2011).

97. Pitstick, A. L. et al. Aggregation of cryopreserved mid-hindgut endoderm for more reliable and reproducible hPSC-derived small intestinal organoid generation. Stem Cell Rep. 17, 1889–1902 (2022).

98. Childs, C. J. et al. EPIREGULIN creates a developmental niche for spatially organized human intestinal enteroids. JCI Insight 8, e165566 (2023).

99. Buenrostro, J. D., Giresi, P. G., Zaba, L. C., Chang, H. Y. & Greenleaf, W. J. Transposition of native chromatin for fast and sensitive epigenomic profiling of open chromatin, DNA-binding proteins and nucleosome position. Nat. Methods 10, 1213– 1218 (2013).

100. Langmead, B. & Salzberg, S. L. Fast gapped-read alignment with Bowtie 2. Nat. Methods 9, 357–359 (2012).

101. Zhang, Y. et al. Model-based Analysis of ChIP-Seq (MACS). Genome Biol. 9, R137 (2008).

102. Pedregosa, F. et al. Scikit-learn: Machine Learning in Python. Mach. Learn. PYTHON.

103. Heinz, S. et al. Simple Combinations of Lineage-Determining Transcription Factors Prime cis-Regulatory Elements Required for Macrophage and B Cell Identities. Mol. Cell 38, 576–589 (2010).

104. Love, M. I., Huber, W. & Anders, S. Moderated estimation of fold change and dispersion for RNA-seq data with DESeq2. Genome Biol. 15, 550 (2014).

105. McLeay, R. C. & Bailey, T. L. RMeseoartchifarEticnlerichment Analysis: a unified framework and an evaluation on ChIP data. (2010).

106. Rauluseviciute, I. et al. JASPAR 2024: 20th anniversary of the open-access database of transcription factor binding profiles. Nucleic Acids Res. 52, D174–D182 (2024).

107. Dobin, A. et al. STAR: ultrafast universal RNA-seq aligner. Bioinformatics 29, 15–21 (2013).

108. Kang, H. M. et al. Multiplexed droplet single-cell RNA-sequencing using natural genetic variation. Nat. Biotechnol. 36, 89–94 (2018).

109. Zheng, G. X. Y. et al. Massively parallel digital transcriptional profiling of single cells. Nat. Commun. 8, 14049 (2017).

110. Wolf, F. A., Angerer, P. & Theis, F. J. SCANPY: large-scale single-cell gene expression data analysis. Genome Biol. 19, 15 (2018).

111. Zhang, Y., Parmigiani, G. & Johnson, W. E. ComBat-seq: batch effect adjustment for RNA-seq count data. NAR Genomics Bioinforma. 2, lqaa078 (2020).

112. Meers, M. P., Tenenbaum, D. & Henikoff, S. Peak calling by Sparse Enrichment Analysis for CUT&RUN chromatin profiling. Epigenetics Chromatin 12, 42 (2019).

113. Krueger, F. & Andrews, S. R. Bismark: a flexible aligner and methylation caller for Bisulfite-Seq applications. Bioinformatics 27, 1571–1572 (2011).

